# Genetic crosses reveal genomic loci responsible for virulence in *Cryptosporidium parvum* infection

**DOI:** 10.1101/2025.05.20.655157

**Authors:** Sebastian Shaw, Xue Li, Gracyn Buenconsejo, Tiffany H. Zhou, Allison Cohen, Daniel Yasur-Landau, Rui Xiao, Daniel P. Beiting, Timothy J.C. Anderson, Boris Striepen

**Affiliations:** Department of Pathobiology, School of Veterinary Medicine, University of Pennsylvania, Philadelphia, Pennsylvania, USA; Texas Biomedical Research Institute, San Antonio, Texas, USA; Division of Parasitology, Kimron Veterinary Institute, Bet Dagan, Israel

## Abstract

The relationship between parasite genotype and pathogenesis is largely unknown for *Cryptosporidium*, a leading cause of diarrheal disease in children. An array of parasites with similar genomes produces varied disease outcomes in different hosts. Here, we isolate and characterize *Cryptosporidium parvum* strains that show marked differences in virulence and persistence in mice. Taking advantage of the sexual lifecycle of this eukaryotic pathogen, we use genetic crosses to discover the underlying chromosomal loci. Whole-genome sequencing and bulk segregant analysis of infection selected progeny mapped three loci on chromosomes 2, 6, and 7 associated with the ability to colonize and persist in mice and the positions of drug resistance genes. The chromosome 6 locus encodes the hyper-polymorphic surface glycoprotein GP60. Reverse genetic studies in both parental strains demonstrate that GP60 controls parasite burden and virulence, but not persistence, and reveal the dominance of the less virulent allele, suggesting it restricts virulence.

## Introduction

The individual outcomes of infectious diseases are highly variable and can range from fatal illness to the absence of notable signs and symptoms. While the basis of this variability is not always understood, multiple and at times interacting contributions of host, pathogen, and environment have been reported. This is also the case for infection with the single celled eukaryotic parasite *Cryptosporidium*. *Cryptosporidium* is a leading cause of diarrheal disease and early childhood morbidity and mortality^1,2^. Infection occurs around the world and under a variety of epidemiological scenarios. Transmission can be linked to drinking or recreational water contact, direct human to human spread, or zoonotic spillover from a variety of domesticated animals. Environmental factors control the endurance and transmissive capacity of the infectious oocyst stage^3^, and abundance or lack of nutrition dramatically impacts the course of infection in children^4–6^. Host genetics^7^, immune status^8^, and co-infection and microbiome^9^ status are similarly important. Individuals who lack T cell function, either from birth or due HIV-AIDS or immunosuppressive treatment are highly susceptible to severe cryptosporidiosis^10^, while humans and animals exposed to repeated infection develop non-sterile immunity that protects them from disease^8,11,12^. Many species and strains of *Cryptosporidium* have been described^13^, and numerous recent reports highlight strain specific (and thus likely pathogen-mediated) differences in host range, transmissibility and virulence^12,14–16^; however, the underlying molecules and mechanisms remain largely unknown.

As typical for apicomplexan parasites, the *Cryptosporidium* lifecycle features phases of asexual and sexual replication, however, in this parasite both phases occur in the same single host. Recent studies have demonstrated that progression from the initial asexual phase to sexual reproduction is obligate, occurs roughly every 48 hours, and that this progression and the resulting sex is essential to continuous infection of the host^17–19^. Remarkably, *Cryptosporidium* will thus experience significant genome recombination and meiotic segregation numerous times over the course of a single infection^20^. We take advantage of this unique sexual lifecycle to associate specific parasite genes with the traits of virulence and persistence through genetic crosses and linkage analysis. Genetic crosses have been transformational to the understanding of drug resistance and virulence in the apicomplexan parasites *Plasmodium* and *Toxoplasma*^21,22^. We recently developed a crossing model for *Cryptosporidium* in mice, where selection for dual drug resistance is used to isolate recombinant progeny to the exclusion of selfing parental types^23^. Crosses were feasible within and between species of diverging host speficity. However, while pairing *C. parvum* and *C. tyzzeri* offered significant phenotypic differences and numerous SNPs for mapping, the overall efficiency of mating was low, resulting in modest genomic recombination^23^. Here we establish a system within the species of *C. parvum* that balances significant genomic and phenotypic diversity with high mating efficiency to build a powerful forward genetic model to disect host parasite interaction.

## Results

### Collection and genome sequencing of *C. parvum* strains

To enable efficient genetic mapping of virulence within the species of *C. parvum,* we collected *C. parvum* strains previously isolated from cattle in the United States and Europe (generously shared with us by Drs. Laurent, Olias, and Krücken), and we returned to a study site in Israel where ovine *C. parvum* had been found and characterized previously^24^. Feces from lambs in Israel were collected and tested for the presence of parasites by PCR. Ocysts were isolated from positive samples, and parasites were amplified in *Ifnγ*^-/-^ mice (see Methods section for further detail, all mice used in this study were of C57BL/6 background). All parasite strains were subjected to whole genome sequencing, followed by allignment to the most recent telomere to telomere assembly of the *C. parvum* Iowa II reference genome^25^, and SNP calling (Figure 1A). Analysis of the diagnositic GP60 locus showed strain KVI to be a IId genotype and all others to be IIa^26^ (Figure S1). The KVI genome showed the largest number of SNPs (5166) when compared to the *C. parvum* Iowa II reference genome (Figure 1A, and 1C show the SNP distribution highlighting areas of high (pink) and low (blue) density). We generated both Illumina short reads and Nanopore long reads, which was then used for *de novo* genome assembly. We found this assembly to be highly syntenic to the recent *C. parvum* telomere to telomere genome^25^ (Figure 1B and S3 and Table S1, note that for KVI we were unable to assenble 5 of the 16 telomeres). Based on the number of distinguishing SNPs and previous reports of phenotypic difference between parasites of IIa and IId background^14^ we chose to pursue Iowa II (Bunchgrass Farms) and KVI as a mating pair to map loci underlying host pathogen interaction.

**Figure 1.**
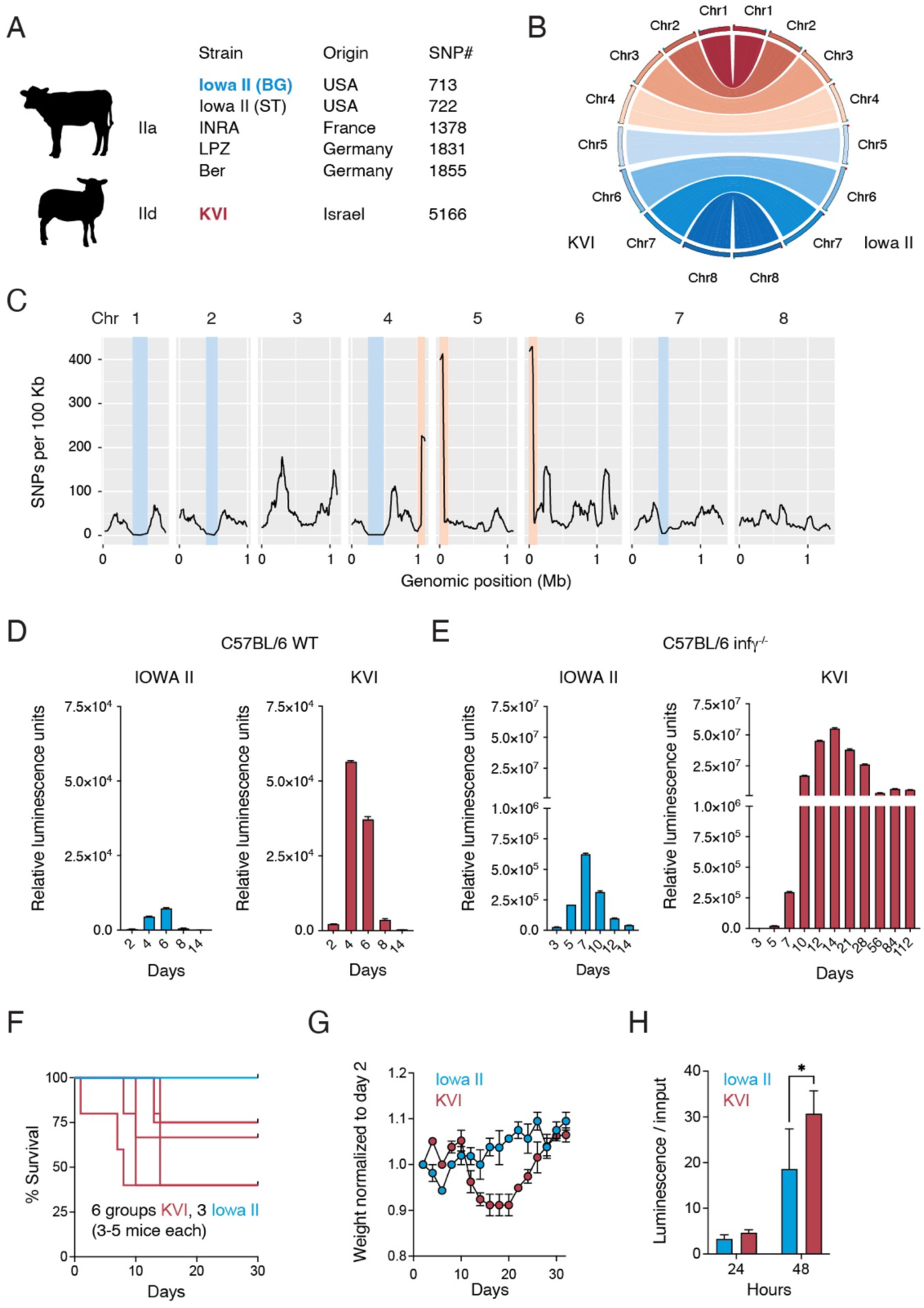
Identification of C. *parvum* strains that differ in virulence and persistence. (A) Type IIa and a newly collected IId strain of *C. parvum* were subjected to whole genome sequencing with KVI showing the largest number of SNPs when compared to the reference genome. (B) The KVI genome was highly syntenic with the reference and (C) shows SNP distribution with areas of especially low (blue) or high SNP density (pink) highlighted. WT (D) or *Ifnγ^-/-^* mice (E) were infected with Iowa II (blue) or KVI (red) parasites expressing Nluc, and parasite burden was measured by following fecal luciferase activity. KVI yields much higher infection and persists (in the absence of IFNγ). (F) Survival of mice infected with the indicated strain, each line shows a cage of 3-5 mice, 6 cages for KVI and 3 for Iowa II. (G) Mice infected with the strain indicated were weighed every two days and weight is plotted normalized to day 2 of infection. Data are mean ± SD of n = 2 biological replicates. (H) HCT-8 cultures were infected with the indicated strain and luminescence was measured after 24 and 48h and plotted normalized to input used for infection (individual cultures are shown as dots and their mean as bars). Data are mean ± SEM of n = 3 biological replicates. Significance was evaluated using 2-way ANOVA, p < 0.05(*).

### Iowa II and KVI show marked differences in virulence and persistence during mouse infection

To facilitate rigorous side-by-side comparison, we engineered the KVI strain to express nanoluciferase (Nluc, Figure S2A) to match Iowa II reporter parasites^23^ already in hand. We integrated a cassette encoding Nluc as well the tdNeonGreen fluorescent protein into the phenylalanyl-tRNA synthase locus selecting for resistance to BRD7929^23^. Age and sex matched C57BL/6 wildtype mice were infected with 20,000 oocysts per mouse of either transgenic Iowa II or KVI, and parasite shedding was measured following luciferase activity in the feces. Both parasite strains produced brief infection of modest burden as typical for *C. parvum* in immunocompetent adult mice^12^, with KVI exceeding the burden of Iowa II (Figure 1D). Next we infected *Infγ*^-/-^ mice with 10,000 oocysts per mouse, which revealed pronounced strain differences. Iowa II resulted in a self-limiting two week course of infection peaking at the level previously reported^12^. By contrast, KVI infection peaked 100-fold higher and remained continuously and robustly patent to the latest time point we measured, day 112 of infection (Figure 1E). 25-50% of *Infγ*^-/-^ mice infected with KVI died by day 14, the peak of parasite burden, while all Iowa II infected mice survived (Figure 1F). This difference in severity was also apparent when following the weight of infected mice (Figure 1G). While Iowa II infected mice showed only a brief and moderate weight loss, those mice infected with KVI lost weight to a nadir at day 14. Mice that survived regained weight afterwards despite continued high infecton. Lastly, we measured growth of both strains *in vitro* using a luciferase assay in human HCT-8 cell cultures^27^. Over 48h of growth KVI was significanty, yet moderately elevated when compated to Iowa II (Figure 1H).

### Genetic cross of Iowa II and KVI parasites

To discover the genes that underlie the difference in mouse infection between the two parasite strains we conducted genetic crosses. *Ifnγ*^-/-^ mice were each infected with 8000 oocysts per mouse of the less virulent Iowa II and 2000 oocysts per mouse of the KVI strain and subjected to dual drug selection with BRD7929 and paromomycin from day 4 to 10 (Figure 2A, Figure S4A-C, cross 1-3). We also conducted crosses in which mice were left untreated (Figure S4-E, cross 4-5). Fecal luciferase activity was recorded to track parasite burden throughout the infections and Figure 2B (drug treated) and 2C (untreated) show measurements for both types of experiments. Parasite burden was slightly higher in experiments that used drug treatment. This likely reflects the effect of the antibiotic paromomycin on the bacterial microbiome which is known to enhance parasite infection^9,27^. We observed sustained infection in all experiments suggesting selection for the superior ability of the KVI parental strain to colonize and persist in mice. We collected oocysts for DNA sequencing across the infection and into the persistence phase and specific time windows for each collection pool are indicated by grey boxes in Figure 2 and S4. Genomic DNA was extracted from 1 to 5 million purified oocysts of each pool. We conducted whole genome Illumina sequencing and variants were called (SNP loci with coverage below 30-fold in either of the compared pools were excluded from analysis).

**Figure 2.**
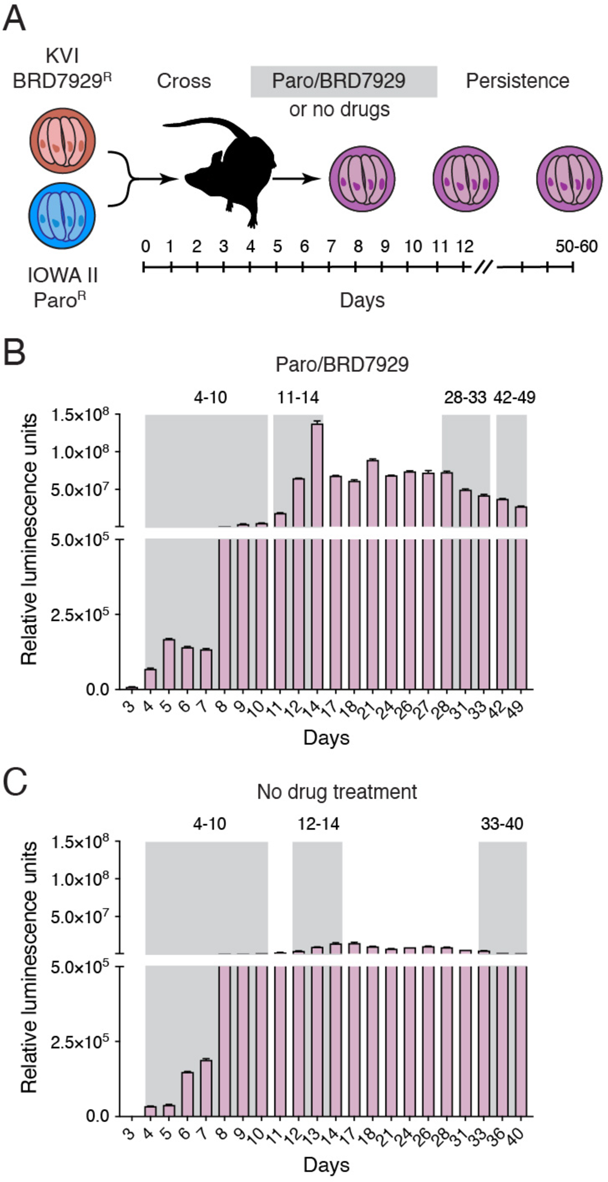
Genetic crosses of Iowa II and KVI parasites. (A) KVI and Iowa II parasites were engineered to be resistant to BRD7929 or paromomycin (paro) respectively. *Ifnγ^-/-^* mice were infected with both strains and treated with both drugs from day 4 to 10, or not. 5 crosses were conducted in total, all are shown in Figure S4). (B, C) Parasite burden was measured by following fecal luciferase activity for two exemplary crosses, one with drug treatment (B) and one without (C). Grey boxes indicate pools of collected feces that were used for oocyst purification and whole genome sequencing.

### Drug resistance loci are readily identified by bulk segregant analysis

Genetic crosses in *Plasmodium* and *Toxoplasma* traditionally relied on the isolation of clonal lines of recombinant progeny and assessed genotype and phenotype individually for each clone to establish linkage^21,22^. Here we turned to bulk segregant analysis as a method that uses progeny populations instead of clones to conduct genetic mapping (for a broader conceptual review of this approach see^28^). The key element of bulk segregant analysis is selection; by comparing allele frequencies across the entire genome among progeny exposed to different selective pressures, we can identify loci for which inheritance from one parent appears enriched or depleted in response to pressure. This is well suited for *Cryptosporidium* where we can collect parasites across the course of in vivo infections with ease but lack a continuous tissue culture system required for cloning.

We first asked whether our crosses might identify the positions of the marker transgenes used. We compared three paromomycin/BRD7929 treated, and two untreated crosses (Figure S4), using the genome sequences of the first oocyst pool collected in each experiment. Visual inspection of the aligned Illumina reads showed loss of the wildtype sequence, and fixation of the resistance genes in the *pheRS* and *TK* loci in all drug-selected pools (Figure S5). Next, we counted reads corresponding to the genotypes of each parent, and calculated allele frequencies for all variable SNPs. These frequencies were then used for bulk segregant analysis. We identified two quantitative trait loci (QTL, Figure 3A) on chromosome three and five, respectively, each characterized by very high G values (G values represent the level of statistical support at each individual SNP, G’ curves average across multiple neighbouring SNPs). Zoomed in views of the allele distribution of these chromosomes show them to be inherited in equal measure from each parent in the absence of drugs (Figure 3B and C, blue shades), but to be highly selected upon drug treatment for the parent that carried the respective marker (Figure 3B and C, red shades). We found the insertion sites at or in close vicinity of the highest confidence SNP. We noted a difference in the shape of the allele frequency and G’-value curve for the markers. While the *TK* locus was found in the middle of the overall peak and the 95^th^ percentile confidence interval, *PheRS* was located outside that interval, and right at the transition between plateau and slope of a broader peak (Figure 3D and 3E). We conclude that *Cryptosporidium* crosses combined with bulk segregant analysis readily identify genes under drug selection with high confidence. The results of the crosses are highly reproducible upon multiple repeats (Figure S6 and S7), and SNP-by-SNP analysis may be superior to reliance on broad, weighted moving averages to identify linked genes.

**Figure 3.**
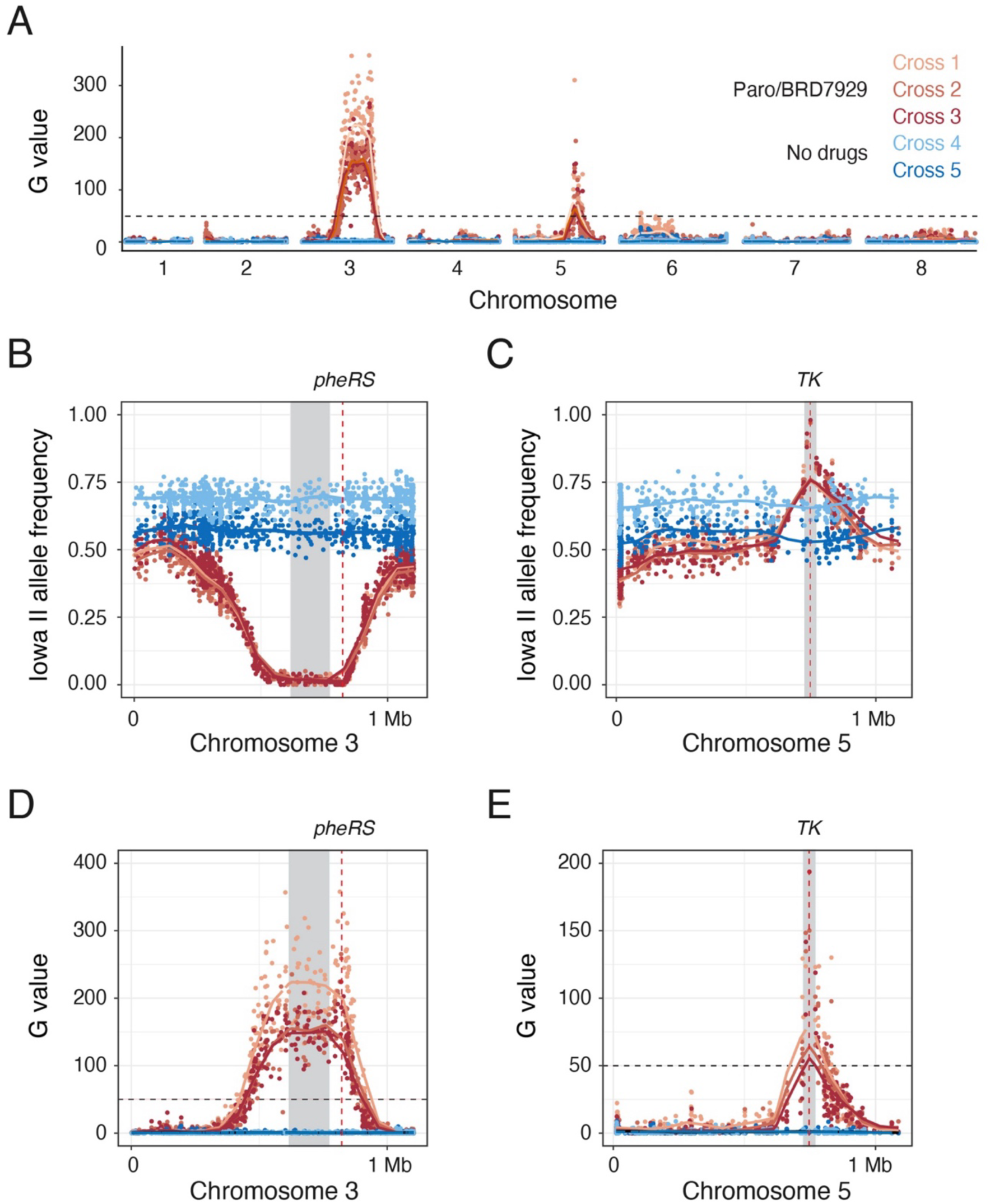
Mapping of drug resistance loci by bulk segregant analysis. (A) Bulk segregant analysis comparing three treated (red) with two untreated (blue) genetic crosses between KVI and Iowa II parasites (the first pool of oocysts collected in each experiment was used for this analysis). The G-value for each individual SNP is shown as dot, the lines are the weighted moving averages for the G’-values, and the significance threshold as a black dashed line. Note highly significant QTL on chromosomes 3 and 5. (B and C) Allele frequences of SNPs using the Iowa II genome as reference for chromosome 3 and 5. While untreated parasites show balanced inheritance from both parents, treatment shifts regions of chromosome 3 to KVI and chromosome 5 to Iowa II respectively. (D and E) Zoomed in chromosomal views G values for both loci. Note that SNPs with peak G values fall into close proximity of the PheRS and TK loci (vertical red dashed lines) which carry the engineered drug resistance determinants. The 95^th^ percentile of each QTL are shown in grey.

## Loci on chromosomes 2,6, and 7 are linked to virulence and/or persistence

We hypothesized that analogous to drug treatment, we might be able to use the host to impose selection suitable to mapping by bulk segregant analysis. We considered that such selection may act immediately, based on differences in the ability to colonize a specific host species, or only unfold over time, reflecting differences in parasite persistence or interactions with the host’s developing adaptive immunity. To account for these scenarios, we followed the progeny through infection and sampled and whole-genome sequenced longitudinally at multiple points of infection for 40-49 days (Supplementary Figure S4). We observed three QTL on chromosome 2, 6, and 7, with marked changes in allele frequency (Figure 4A and 4B, Figure S7 and Table S1 for an overview of G- and G’-values of all five crosses). These changes occurred regardless of whether parasites were subjected to drug treatment or not, and all three reflected preferential inheritance of the KVI genome. Bulk segregant analysis identified highly significant QTL for these positions, however there were kinetic differences. While the locus on chromosome 6 showed almost immediate impact and reached significance at 11-14 days, the two other loci produced shifts in allele frequences and significant QTLs only after the infections proceeded to later time points (Figure 4, C-D).

**Figure 4.**
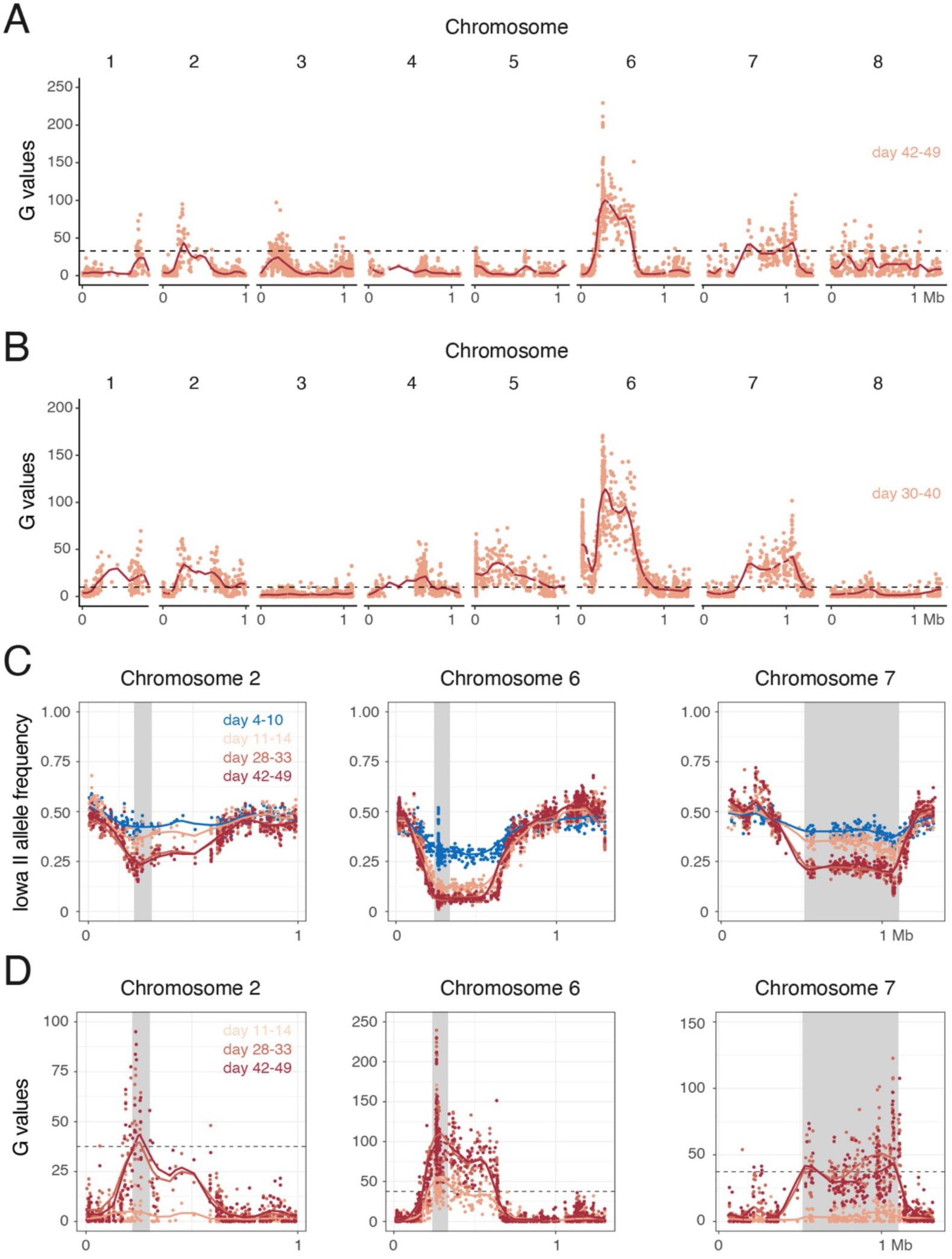
Longitudinal analysis of genetic crosses identifies loci associated with virulence and persistence. Genetic crosses were conducted between Iowa II and KVI parasites with (A) or without (B) treatment with BRD7929 and paromomycin. Parasites were sampled as shown in Figure 2D and C and whole genome sequenced. G-values of bulk segregant analysis are shown comparing the first and last sampling window for each experiment. There are three significant QTL in both treated, and untreated populations. (C) Allele frequencies and (D) G values for these three loci shown in more detail for the cross that used drug selection (all experiments are found in Figure S6). While the QTL on chromosome 6 emerges early, the loci on chromosomes 2 and 7 only show selection upon prolonged infection (samples from day 26 onwards). In all plots the horizontal dashed line represents the threshold for QTL detection. The 95^th^ percentile of each QTL are shown in grey.

### The locus on chromosome 7 contains dense granule proteins, one of which is required for high parasite burden

The QTL peak on chromosome 7 encompasses multiple genes including three presumptive secretory proteins and two ABC transporters. Measurements of mRNA abundance comparing the two parental strains found no significant differences for any of these genes (Figure 5A). Three genes cgd7_4490, cgd_4500, and cgd_4530 were predicted to likely encode dense granule proteins in a recent spatial proteomic study^29^ with cgd7_4500 showing the highest degree of polymorphism and a dN/dS ratio of 2.93 across the sequences currently available on the CryptoDB database. We began analysis by introducing sequences encoding a C-terminal influenza hemagglutinin (HA) epitope into cgd7_4490 and cgd_4500 in the KVI strain background (Figure S2B). Immunofluorescence assays using anti-HA antibodies using expansion microscopy confirmed that both proteins localize to the dense granule compartment in sporozoites and we will thus refer to the proteins as dense granule protein DG5 and DG6, respectively (Figure 5B and 5C). Following secretion during invasion, DG5 is found in the parasitophorous vacuole surrounding the intracellular parasite, while DG6 localized to the host parasite interface. Next, we targeted each locus in the KVI strain by introducing a Nluc/Neo marker that disrupted the coding sequence (Figure S2B). Drug resistant parasites were recovered in both cases, albeit with pronounced delay for DG6. Uniform disruption was confirmed by PCR mapping of the loci (Figure S2B). In side-by-side infection experiments in *Infγ^-/-^* mice we noted a ∼30-fold reduction of peak parasite burden for the ΔDG6 mutant, while deletion of DG5 had comparatively little impact on infection (Figure 5D and 5E). Consistent with reduced parasite burden, mice infected with ΔDG6 mutants showed less weight loss when compared to those infected with KVI wild type (Figure 5F). In HCT-8 tissue culture experiments *in vitro* growth of the ΔDG6 mutant was found to be indistinguishable from KVI wild type (Figure 5G). To rigorously control whether the observed burden reduction was due to modification of the target locus alone, we next complemented the mutant. We reintroduced the gene amplified from either the Iowa II or the KVI strain as a transgene 3’ to the PheRS locus using BRD7929 selection, including the respective 5’ and 3’ sequences from each allele type to ensure native levels of expression (Figure S2C). Parasite burden in both complemented strains rebounded to levels comparable to KVI wild type while the ΔDG6 mutant again showed lower infection (Figure 5H).

**Figure 5.**
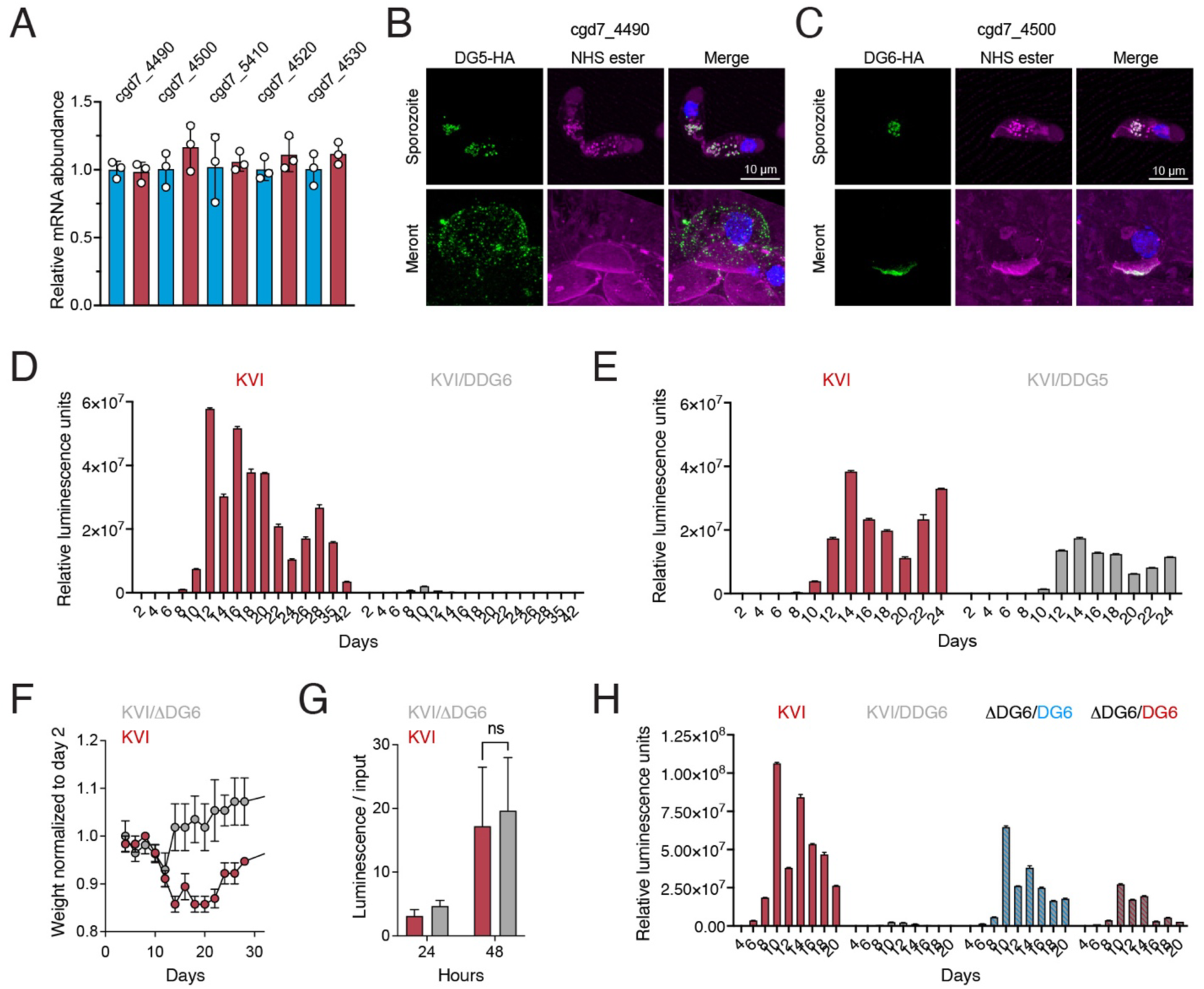
Loss of dense granule protein DG6 but not DG5 results in attenuation in vivo. (A) The core of the QTL on chromosome 7 includes 3 presumptive dense granule proteins and 2 ABC transporters. Comparison of their relative expression level by qPCR found no difference between KVI an Iowa II (a representative of two experiments with three biological and 3 technical replicates each is shown). The ΔΔCt method was used to determine the relative expression with GAPDH as the control gene. The fold change of the KVI strain is relative to GAPDH control and Iowa II samples. Data represents three biological repeats and is plotted as the mean. A list of all the primers is provided in Supplementary Table S2. (B and C) Expansion microscopy fluorescence micrographs of HA tagged DG5 (B) and DG6 (C) in green, NH-ester in magenta and Hoechst in blue. (D and E) *Ifnγ^-/-^* mice were infected with ΔDG5 (D) or ΔDG6 (E) mutant parasites in direct comparison with the KVI parental wild type and parasite burden was measure by fecal luciferase assay. (F) Body weight of mice infected with ΔDG6 or KVI parent normalized to the weight at day 2 of infection. (G) HCT-8 cell culture assays comparing growth of ΔDG5 and ΔDG6 to KVI parent. (H) The ΔDG6 was complemented in trans with the DG6 gene of Iowa II (blue) or KVI (maroon). These parasites were used to infect *Ifnγ^-/-^* mice in comparison to the parental ΔDG6 and KVI wild type. Both genes confer enhance infectivity to the mutant.

We conclude that while DG5 is dispensable under the conditions tested, loss of DG6 results in reduced infection *in vivo* but not *in vitro*, and this is rescued by complementation. We observed that sequences from either parent provided similar levels of complementation, suggesting the strain specific difference responsible for the QTL to reside within a different gene.

## GP60 controls parasite burden and virulence but not persistence

Figure 6A shows a closer inspection of the QTL on chromosome 6, plotting average SNP G-values for bins of 1000 bp, along with annotated gene models (all individual G-values are found in Table S1). The peak falls into gene *cgd6_1080* which encodes glycoprotein (GP) 60, a polymorphic gene that is frequently used to type *Cryptosporidium*^26,30^. GP60 has been suspected to be linked to virulence^31^, and a recent study has explored its contributions to host infectivity^32^. When comparing the GP60 gene between the parental Iowa II and KVI strains we noted 149 SNPs and 3 indels (Figure S8A), and found that the selected progeny largely carries the KVI allele (Figure 4C). Both alleles encode an N-terminal signal peptide, a C-terminal glycosyl-phosphatidylinositol anchor signal, numerous potential O-glycosylation sites, and a single potential N-linked glycosylation site, albeit not at the same position in the protein (Figure S8B shows an annotated sequence alignment). The QTL includes 5’ upstream regions with the presumptive promoter, and we thus considered that KVI and Iowa II might express GP60 at different levels as previously noted for ROP18 alleles in *T. gondii* strains that differ in virulence^33,34^. However, qPCR analysis of both parents showed similar GP60 mRNA abundance (Figure 6B). Next, we used reverse genetic engineering to test the impact of GP60 in isolation. We considered that GP60 from the more virulent KVI strain might be a dominant factor conferring virulence. We introduced KVI GP60 along with its 5’ and 3’ regulatory sequences as a transgene into the dispensable thymidine kinase locus of Iowa II parasites (this also introduced a Nluc reporter, Figure 6C, Figure S2D). When we infected *Infγ^-/-^* mice with the resulting stable transgenic parasites we found, to our surprise, the resulting infection to be moderate and similar to Iowa II (Figure 6E). Hypothesizing that alternatively, the low virulence Iowa II allele might be dominant, we next disrupted the coding sequence of the endogenous GP60 gene by insertion of a PheRS marker cassette in the bi-allelic strain (Figure 6D, Figure S2D). This resulted in a marked increase in parasite burden in side-by-side infections (Figure 6E). We also engineered reciprocal modifications in the highly virulent KVI parent (Figure S2E). Expressing Iowa II GP60 as a transgene decreased KVI parasite burden, again suggesting the dominance of this allele in reducing virulence (Figure 6F). We also considered that expression of two distinct alleles of GP60 might have a dominant negative effect limiting parasite growth in a way unrelated to virulence. To control for this, we ablated the original KVI gene in the pseudo diploid strain leaving only the single Iowa II allele behind. This did not change parasite burden, which remained low, arguing that parasite growth *in vivo* is not impeded by the expression of two alleles of GP60 but by the presence of the Iowa II allele (Figure 6F). We monitored the weight of all infected mice as a measure of virulence. Removing Iowa II GP60 in Iowa II alone was not sufficient to cause enhanced weight loss, and this might be linked to the fact that while parasite burden increased dramatically the duration of the infection remained short (Figure 6G). Expression of Iowa II GP60 prevented the substantial weight loss associated with KVI infection (Figure 6H). When these replacement mutants were used to infect HCT-8 cultures we observed that replacing GP60 in the Iowa II strain with the KVI allele enhanced growth, while exchange of GP60 in the KVI strain reduced growth when compared to the respective parent (Figure 6I and J). We were unable to ablate GP60 completely in the strains we studied. We conclude that bulk segregant analysis correctly mapped and identified GP60 as a major strain-specific contributor to virulence. Interestingly, reverse genetic analysis in both parental backgrounds suggests that Iowa II GP60 moderates virulence in a dominant fashion. While changes in GP60 changed virulence they had no impact on persistence, Iowa II did not gain persistence, and KVI did not lose it when the gene was replaced.

**Figure 6.**
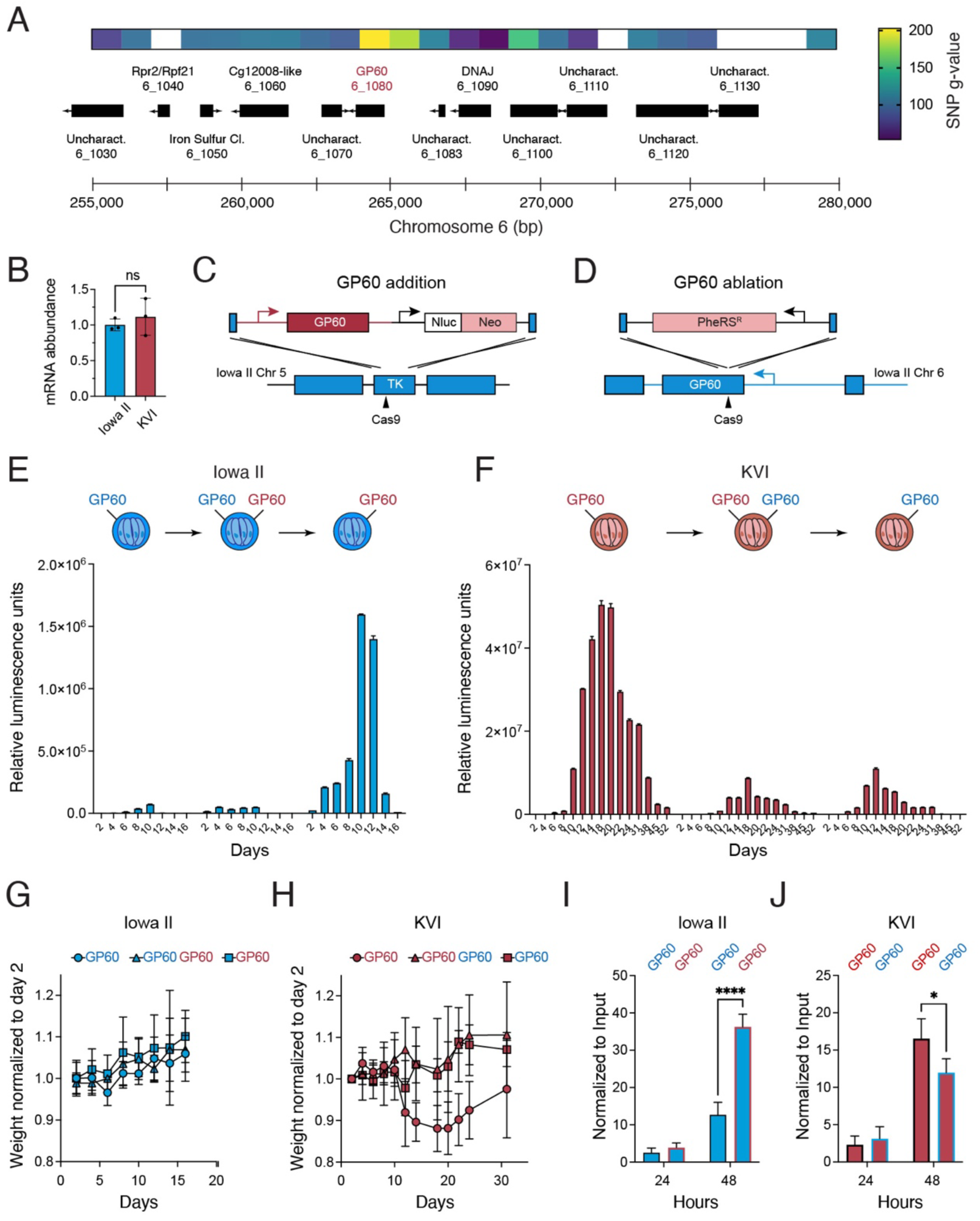
GP60 controls parasite burden and virulence. (A) Section of chromosome 6 that includes the QTL showing the coding regions of genes as black boxes with arrows indicating direction of transcription, and mean G values binned in 1000 bp intervals (white indicates areas devoid of SNPs distinguishing the parents). (B) qPCR analysis of KVI GP60 mRNA abundance relative to IowaII. The ΔΔCt method was used to determine the relative expression with GAPDH as the control gene. The fold change of the KVI strain is relative to GAPDH control and Iowa II samples. Data represents three biological repeats and is plotted as the mean. A list of all the primers is provided in Supplementary Table S2. (C) The GP60 gene including its regulatory regions of the KVI strain was introduced as a transgene in the thymidine kinase (TK) locus of the Iowa II strain using Neo, the native GP60 locus was then disrupted (D) using the PheRS marker generating the strains used in (E). Reciprocal modifications were engineered into KVI (F, see supplementary Figure S2 B and C for detail on all allelic replacement mutants). (E and F) WT parasites, parasite expressing GP60 of both parents, and parasite expressing only the GP60 of the other parent were used to infect *Ifnγ-/-* mice and parasite burden was measured by fecal luciferase assay (note that all strains carried a single Nluc gene). Expression of Iowa II GP60 is associated with lowered burden in both parental backgrounds, while adding or removing KVI GP60 had little effect. (G and H) Mean weight of infected mice from (D and E) normalized to day 2 of infection (combined data of two experiments using four mice each, error bars show standard deviation). (I and J) HCT-8 cultures were infected with the indicated strains and luminescence was measured after 24 and 48h and plotted normalized to input used for infection (individual cultures are shown as dots and their mean as bars). Data are mean ± SEM of n = 3 biological replicates. Significance was evaluated using 2-way ANOVA; p < 0.05(*), p < 0.0001(****).

## Discussion

Sexual recombination has been hypothesized to be critical for the ability of *Cryptosporidium* to adapt to opportunity and challenge through rapid evolution of complex traits. Recent studies have highlighted the likely role of sex in the emergence of enhanced transmissibility, resulting in an epidemiological sweep of *C. hominis* strain IfA12G1R5 across the United States^15,35^. Recombination was similarly documented in studies of *C. hominis* across the African continent^36^. Increased host specialization resulting in narrowing adaptation to humans has been noted in the typically more broadly zoonotic *C. parvum*, and this has been discussed as the potential result of sex^37^. Interestingly, the overall rate of such genomic exchange appears higher in recent human history, likely reflecting increased long-distance travel of both humans and domestic animals enhancing mating opportunities of geographically separated parasite strains^38^. In this study we show through controlled laboratory experimentation that sex indeed allows parasites to recombine and rapidly select beneficial traits including drug resistance and superior colonization ability. We also show that this can be exploited to discover parasite gene function in forward genetic mapping experiments.

While *Cryptosporidium* is not amenable to mapping based on the isolation and study of individual progeny clones, it proves well suited for bulk segregant analysis, which identified the loci of the drug resistance markers used with high statistical support. We noted relatively broad G’ linkage peaks that contain one or multiple much more narrow peaks when considering the G-values of individual SNPs. This could indicate multiple loci to contribute to the overall QTL, however, this was also observed for the PheRS drug resistance locus where a single gene drives selection. For most organisms, crosses involve a single round of recombination, and the progeny thus represent the F1 generation. However, *Cryptosporidium* experiences continuous mating among the progeny as sex occurs every third day of an ongoing infection^17,18^. We note that mapping was successful even in the absence of dual drug selection to enforce mating. This suggest an overall high rate of recombination and that crosses with unmarked strains, e.g. for species not amenable to transfection, are likely feasible. Using marker genes biases selection for their loci, this may fix chromosomes limiting resolution. By the same token, careful placement of markers could enhance segregation of loci difficult to resolve.

Several additional factors likely influence the resolution of genetic mapping including the specific biology of the trait under selection, the number of recombination events between the strains used, and the chromosomal location of the genes to be distinguished. One obvious factor is the number of SNPs that discern the parents. Higher SNP density improves the ability to pinpoint the position of selected genes, as seen in the highly polymorphic GP60 locus (Figure 6), while low density diminishes precision. The remarkable similarity of *C. parvum* IIa strains even when isolated from different continents (Figure 1A) thus was an early impediment to our efforts. IIa strains are highly prevalent in cattle, and parasite uniformity may reflect host uniformity due to the global distribution and dominance of certain highly productive cattle breeds^39^. Isolation of IId parasites from lambs was important to success, and other studies have noted more pronounced differences between IIa and IId strains as well^14,40^. Future sampling of diverse hosts could yield diverse parasites and further improve mapping resolution and provide insight into the mechanistic basis of host specificity. Another factor to consider is the strength of the selection applied^41^. The highly potent BRD7929 kills parasites fast^42^ and the rigorous selection imposed may have resulted in rapid fixation of a large chromosomal segment, while the less potent drug paromomycin provided slower selection, and thus added opportunity for recombination and resolution. Lastly, time itself is important, and the chronic nature of KVI infection in the mouse model chosen here offered the opportunity for longitudinal evaluation. Two of the three QTLs only rose to significance over time, suggesting that they are required for chronic persistence but not initial colonisation.

More work is needed to understand the specific mechanisms by which these loci impact persistence. However, we note the presence of polymorphic secretory proteins, in particular dense granule proteins, a subset of proteins that in *T. gondii* was shown to contribute to immune evasion by manipulating host defense pathways and their transcriptional and regulatory circuits^43–45^. The late emergence of these loci might suggest their dependence on selection imposed by the development of adaptive immunity. Crosses in hosts with varied immune competence may provide differentiating selection to link loci to the evasion of specific restriction mechanisms or may yield additional loci only revealed by the change of selection applied. The advanced tools accumulated for the study of immunity in mice offer a multitude of genetic backgrounds and reagents to remove or add specific elements of the immune system. We and others have shown *C. parvum* to be highly sensitive to such manipulations (reviewed in ^10,11,46^). This breadth of selection systems combined with the relative ease of the population-based crossing model developed in this study holds significant promise for future of forward genetic analysis of host pathogen interaction in *Cryptosporidium*.

GP60 stands out as the most significantly selected locus in our experiments leading to rapid fixation of the KVI allele indicating that it exerts a profound influence on the ability of *C. parvum* to colonize mice, reach high parasite burdens, and to cause disease as measured by weight loss and mortality. Reverse genetic analysis rigorously validated this finding – replacing the GP60 locus alone changes parasite load and virulence in both parental genomes as background. The well acknowledged high degree of polymorphism of this locus^13,26^ suggests that it is under significant diversifying selection^47^, a hallmark of rapid host-pathogen coevolution. GP60 is proteolytically processed into GP40 and 15 which remain associated following cleavage and membrane bound due to a GPI anchor on GP15^48^. GP40 is particularly variable and heavily glycosylated^30,49^ (Figure S8). The protein has been localized to the surface of extra and intracellular parasite stages, and this localization depends on the presence of signal peptides for ER insertion and GPI anchor addition^32^. GP60 has been suggested to play a role in host cell invasion^49^, however, its specific molecular function remains to be revealed. Li and colleagues found that replacing GP60 in a IId strain with a IIa allele resulted in less virulent parasites^32^. We engineered replacement mutants in IIa and IId parental backgrounds along with intermediate pseudo-diploid strains expressing both alleles. Phenotypic analysis of these mutants demonstrated dominance of the less virulent Iowa IIa allele, in both parental backgrounds virulence was restricted by the presence of the Iowa II allele, rather than the KVI allele enhancing virulence. This is intriguing, and the mechanism underlying this dominance is still unknown. Different GP60 allele might impact on the parasite’s ability to invade and grow. While the strain difference was noticeable in tissue culture, the effect was much more pronounced in animals, particularly in those lacking IFNγ. Overall, this may suggest that GP60 is recognized by the host as a trigger for immune restriction, and that the IIa allele produce a stronger response in mice. Such a scenario underlies the relative burden of *T. gondii* in different rat hosts. Dense granule proteins secreted by the parasite trigger a potent inflammasome response restricting the pathogen^50^. *C. parvum* activates a range of innate mechanisms through the NLRP6 inflammasome^51,52^, a subset of toll like receptors^53,54^, and likely additional pathways^55^. GP60 is also a target of adaptive immunity and a prominent antigen recognized by antibody and cell mediated responses^49,56,57^. Genetic mapping studies in *Plasmodium* have identified strain specific antigens^58^. Exploring the impact of GP60 alleles in the context of mouse mutants lacking specific restriction pathways will be helpful to further distinguish these models.

The crosses reveal the amplitude of parasite burden and the persistence of parasite presence as distinct strain specific traits governing *Cryptosporidium* infection. GP60 controls parasite burden in the mouse model but it does not impact on persistence. Switching the GP60 allele to KVI in the Iowa parent, parasites did not gain persistence, nor did the KVI parent lose persistence in the reverse swap. Additional loci on chromosomes 2 and 7 were significantly selected in chronic infection suggesting that they may harbour persistence conferring genes. Initial studies on two genes encoding dense granule proteins demonstrated one of them, DG6 to be important for high parasite burden *in vivo* but not *in vitro*. However, complementation studies argue against a strain specific function of this factor, making additional genes in this locus high value candidates to be pursued in future studies. Importantly, both high burden and longer persistence were required for the KVI strain to cause severe life-threatening disease.

Our experimentation shows that forward genetic studies in *Cryptosporidium* are feasible, and with the help of bulk segregant analysis highly capable to discover the genomic location of genes associated with drug resistance and virulence. Continued recombination and continued selection yields linkage of different parasite genes to different phases of the infection. Varying the specific circumstance under which host, and parasite encounter each other will tune the selection pressure, and is likely to reveal different aspects of interaction and the numerous parasite factors that modulate such interaction. The small genome size of *Cryptosporidium* and the ability to use mice to conduct crosses makes this a straightforward discovery model to understand how host restriction and parasite evasion govern parasite infection and the evolution of host specificity.

## Acknowledgements

This work was supported in part by grants from the National Institutes of Health to B.S. (R01AI112427, R01AI127798), B.S. and Christopher Hunter (R01AI148249), T.J.C.A. (P01AI127338, R01AI048071), and Swiss National Science Foundation fellowships P2BEP3_191774 and P500PB_211097 to S.S. and the NIH fellowship F32 AI183654 to A.C. We thank Jessica Byerly and Chloe Tang for help with animal experiments, the PennVet imaging facility for microscopy support, Daniel J. Cutillo and Lisa Mattei for help with DNA sequencing, and Sebastian Lourido for critical reading.

## Author Contributions

S.S. and B.S. conceived the study. D.Y.L. isolated and genotyped the KVI strain in Israel. S.S. performed genetic crosses with support from G.B., and reverse genetic characterization experiments with contributions from T.H.Z. to tagging and gene ablation, and A.C. to expansion microscopy. X.L. performed bulk segregant analysis with support from T.J.C.A. and R.X. generated the automated bulk segregant analysis and the de novo assembly of the KVI strain genome. S.S. and B.S. wrote the manuscript with contributions from all authors.

## Resource availability

Lead contact

For access to reagents or parasite strains used in this study please contact the lead contact, Boris Striepen.

## Materials availability

All unique/stable reagents generated in this study are available from the lead contact with a completed Materials Transfer Agreement.

### Experimental model and subject details

#### Mouse models of infection

All parasites were maintained in 6-to 8-wk-old male and female C57BL/6 *Ifnγ^−/−^* (Jackson Laboratory stock no. 002287) mice bred in-house. C57BL/6 wildtype mice were purchased from Jackson. Protocols for animal maintenance, care and experimentation were approved by the Institutional Animal Care and Use Committee of the University of Pennsylvania (protocol #806292).

#### Parasite strains

Oocysts of *C. parvum* strains were obtained originally from the following sources: Iowa II (BG), Bunch Grass Farms, Deary, Idaho; Iowa II (ST), Dr. Reed, University of Arizona; LPZ and Ber, Dr. Olias, University of Giessen, Germany, and Dr. Kücken, Free University Berlin, Germany; INRAE^59^, Dr. Fabrice Laurent, INRAE and University of Tours, Nouzilly, France. KVI strain oocysts were obtained in March 2023 from a clinical case of cryptosporidiosis in a 2.5-week-old lamb on a farm in Beit Dagan, Israel. Fecal samples found to harbor oocysts detected using modfied Ziehl-Neelsen staining were subjected to DNA extraction using the Presto Stool DNA Extraction Kit (Geneaid, Taiwan). The species of *Cryptosporidium* was determined by sequencing PCR amplified SSU rRNA^60,61^ and subtyped using GP60 amplicons^62^. Amplicons were matched to sequences deposited in GenBank using BLAST^63^, and GP60 subtype was established by multiple sequence alignment with IIa and IId subtypes and inspection^64^. For isolation, oocysts were subjected to sedimentation as described in the WOAH Terrestrial Manual^65^ with the omission of ethyl acetate, and all centrifugation steps were performed for 10 min at 2000 RPM and 4°C. Recovered oocysts were washed once in 0.5% household bleach prior to wash, storage and shipment in PBS at 4°C.

#### Cell culture

HCT8 obtained from ATCC were cultured in DMEM supplemented with 10% FBS at 37°C in presence of 5% CO2.

## Method Details

### Generation of transgenic parasites

To generate transgenic parasites, 1.25 × 10^7^ oocysts were incubated at 37 °C for 1 h in 10 mM HCl followed by two washes with phosphate buffered saline (PBS) and an incubation at 37 °C for 1 h in 0.2 mM sodium taurocholate and 22 mM sodium bicarbonate to induce excystation^66^. Excysted sporozoites were electroporated and used to infect four- to eight-week-old mice pretreated with antibiotics by oral gavage as previously described^27,67^. Transgenic parasites were selected with paromomycin in the drinking water (16g/L) or BRD7929 by daily gavage (5mg/kg/day formulated in in 70% PEG400 and 30% aqueous glucose (5% in H_2_O). Feces were collected from each mouse in the cage, mixed, and 20 mg were used to measure parasite burden by nLuc luciferase assay^27,67^. All transgenic parasites generated in this study are documented in Figure S2 and in Star Methods Table S2.

### KVI tdNeon

The repair encodes the last 113 bp of the pheRS gene (cgd3_3320, recodonized) including the mutation that confers resistance (L482V) to BRD7929. This short sequence is followed by a nanoluciferase reporter fused to tdNeonGreen under the enolase promoter.

### Iowa with ectopic GP60-KVI

The repair encodes the GP60-KVI version under its own promoter and 3’UTR followed by a nanoluciferase reporter fused to the Neo resistance gene under the enolase promoter. It is integrated into the thymidine kinase locus.

### Iowa with ectopic GP60-KVI and endogenous GP60 knockout

The repair encodes the pheRS gene including the mutation that confers resistance (L482V) to BRD7929 followed by a nanoluciferase reporter under the enolase promoter. The parental strain for this transgenic is the Iowa with ectopic GP60-KVI strain.

### KVI with ectopic GP60-Iowa

The repair encodes the GP60-Iowa version under its own promoter and 3’UTR followed by a nanoluciferase reporter fused to the Neo resistance gene under the enolase promoter. It is integrated into the thymidine kinase locus.

### KVI with ectopic GP60-Iowa and endogenous GP60KO

The repair encodes the pheRS gene including the mutation that confers resistance (L482V) to BRD7929 followed by a nanoluciferase reporter under the enolase promoter. The parental strain for this transgenic is the KVI with ectopic GP60-Iowa strain.

### KVI cgd7_4490 knockout

The repair encodes a nanoluciferase reporter fused to the Neo resistance gene under the enolase promoter.

### KVI cgd7_4490 Ha-tag

The repair encodes the HA-tag followed by a nanoluciferase reporter fused to the Neo resistance gene under the enolase promoter.

### KVI cgd7_4500 knockout

The repair encodes a nanoluciferase reporter fused to the Neo resistance gene under the enolase promoter.

### KVI cgd7_4500 Ha-tag

The repair encodes the HA-tag followed by a nanoluciferase reporter fused to the Neo resistance gene under the enolase promoter.

### KVI cgd7_4500 knockout and endogenous Iowa II cgd7_4500

The repair encodes the last 113 bp of the pheRS gene (cgd3_3320, recodonized) including the mutation that confers resistance (L482V) to BRD7929. This short sequence is followed by the Iowa II cgd7_4500 version under its own promoter and 3’UTR. The parental strain for this transgenic is KVI cgd7_4500 knockout strain.

### KVI cgd7_4500 knockout and endogenous Iowa II cgd7_4500

The repair encodes the last 113 bp of the pheRS gene (cgd3_3320, recodonized) including the mutation that confers resistance (L482V) to BRD7929. This short sequence is followed by the KVI cgd7_4500 version under its own promoter and 3’UTR. The parental strain for this transgenic is KVI cgd7_4500 knockout strain.

### Cell Culture and in vitro growth assay

HCT-8 cells were purchased from ATCC (CCL-224TM) and maintained in RPMI 1640 medium (Sigma-Aldrich) supplemented with 10% Cosmic calf serum (HyClone). Cells were infected with 40000 excysted sporozoites and serum was reduced to 1%. Medium was aspirated after 24h and 48 h, cells were lysed and mixed with NanoGlo substrate (Promega) and luminescence was measured using a Glomax reader (Promega). For normalization, luminescence was measured after 4h post infection (input).

### Microscopy

Infected cells were fixed with 4% paraformaldehyde (Electron Microscopy Science) in PBS and stained with Hoechst (1 μg/mL). Coverslips were then mounted on glass slides with fluorogel (Electron Microscopy Science) mounting medium. Ultrastructure expansion microscopy was performed as previously described^29,68,69^. Sporozoites were allowed to excyst in culture media for 2 hours before allowing them to settle onto poly-D-lysine-coated coverslips prior to fixation. For intracellular stages cells were fixed 24 hours post-infection. Crosslinking prevention and anchoring were performed in a mixture of 2% acrylamide and 1.4% formaldehyde overnight at 37°C. Samples were embedded in a hydrogel, which was subjected to denaturation, expansion, and staining with antibodies to detect HA epitope-tagged proteins, Atto647-conjugated NHS ester dyes to detect protein density, and Hoechst to detect DNA. Stained gels were imaged on a Leica Stellaris 8 inverted confocal microscope with a 63X water immersion objective.

### qPCR

Infections were set up in a 24-well plate, with 200,000 oocysts per well. RNA was extracted at 48 h by direct lysis in the well with 350 μl of RLT lysis buffer from the RNeasy minikit (Qiagen). The manufacturer’s instructions were followed, and RNA was eluted in 40 μl of RNase-free water. cDNA was prepared per sample using SuperScript IV reverse transcriptase (Thermo Fisher Scientific). The cDNA was diluted 1:10 before setting up the quantitative PCR (qPCR) in a 10 μl reaction, which included SsoAdvanced Universal SYBR Green Supermix (Bio-Rad). The reaction was loaded into the ViiA 7 Real Time PCR system (Thermo Fisher Scientific) with the following conditions: initial incubation at 95 °C for 3 min, 95 °C for 15 s and 60 °C for 30 s for 40 cycles, and a single melt curve. The ΔΔ*C*t method was used to determine the relative expression with GAPDH as the control gene. A list of all of the primers is provided in Supplementary Table S2.

### Genetic crosses and sample collection

In total, five crosses between the *C. parvum* Iowa II (BG) and KVI strains were conducted. 3 crosses with initial dual drug selection (paromomycin and BRD7929). Mice were infected with 2000/8000 (KVI/BG; cross 1 and cross 2) and 1000/4000 (KVI/BG; cross 3) oocysts per mouse, respectively. Recombinant progeny was selected by dual drug selection from day 4 to day 10. 2 crosses were conducted in the absence of drug selection with 100/9900 (KVI/BG) oocysts per mouse for each cross. Fecal material was collected daily and stored at 4°C. 14 Segregant pools were collected as indicated in Figure S4 (grey boxes).

### Oocyst purification and genomic DNA extraction

Fecal material of infected mice was collected and oocyst were purified using a sucrose gradient and CsCl flotation^67^. Genomic DNA was extracted using phenol/chloroform as described^23^.

### Library preparation and sequencing of cross progeny

We prepared Illumina libraries from extracted genomic DNA and sequenced both parents and 14 segregant pools. The library preparation was carried out using Illumina DNA Prep (former Nextera DNA Flex kit, Illumina Inc.). Subsequently, sequencing was performed on the Illumina NextSeq 2000 sequencer, utilizing P2 300 cycle flowcell kit. We prepared Oxford Nanopore library from whole genome amplified genomic DNA using REPLI-G MINI kit WGA (CAT #150023, Qiagen, Hilden, Germany) followed by a T7 Endonuclease I debranching (CAT #M0302, New England BioLabs, Ipswich, MA) as described^70^. We used native barcoding kit SQK-NBD114.24 to construct ligation sequencing library. Total of 400ng of final library was loaded onto a R10.4.1 flowcell, and sequenced on an Oxford Nanopore MinION machine for 72 hours.

### *De novo* genome assembly

Illumina short read (Nextseq 2000, 2x150bp) and Nanopore long read (MinION R10.4.1) sequences were generated and used for genome assembly. Illumina short reads were adapter and quality (>Q30) trimmed with TrimGalore (v0.6.10). Raw nanopore reads are base called and demultiplexed with ONT Dorado (v0.9.1) with Super Accuracy (SUP) plus duplex mode. CADECT (v1.0.2) was used on base called long reads to remove concatemers resulted from whole genome amplification. Long reads were assembled with Flye (v2.9.5) into 69 contigs. Hybrid genome assembly was performed with SPAdes (v4.0.0) using both illumina short-read and Flye long-read assembled contigs. NCBI fcs-adapter (v0.5.0) and fcs-gx(v0.5.0) were used to remove adapter contamination and non-*Cryptosporidium* sequences from the assembled genome. The genome was re-orientated and scaffolded based on the T2TCpBGF genome ^25^ with RagTag(v2.1.0) into 8 chromosomes. The genome was polished with 3 rounds of illumina short-reads and 3 rounds of Nanopore long-reads, both using Racon (v1.4.3). The final genome size is 9,083,795bp and we identified 11 out of 16 telomeres from the assembly using FindTelomeres. The consistency between the KVI *de novo* assembly and the T2TCpBGF genome was assessed using MUMmer (Figure S3). The Iowa II and KVI genomes aligned well, showing high similarity with only a few small inversions across the genome. A single translocation (∼500bp) was observed at the end of chr 7, caused by rearrangement of the Hemogen gene. The inversions ranged from 100 bp to 53 kb, which are unlikely to impact recombination. We annotated the KVI genome with Liftoff (v1.6.3) using the T2TCpBGF annotation as well as *de novo* BRAKER (v3.0.8). Annotations from Liftoff and BRAKER3 were merged and cleaned with AGAT (v1.4.0). The final annotation of KVI identified 3928 protein coding, 45 tRNA, and 15 rRNA genes.

Reference KVI genome can be accessed from NCBI PRJNA1263880. Alternatively, KVI genome is fasta format and annotation in gff format can also be accessed here: https://github.com/ruicatxiao/Automated_Bulk_Segregant_Analysis/tree/main/KVI_genome_annotation

### Genotype calling

The Iowa II *de novo* assembly (T2T reference genome^25^) was then used as the reference genome to identify single-nucleotide polymorphism (SNP) distinguishing the two parents, which were then used for bulk segregant analysis. Whole-genome sequencing reads for each library were individually mapped to the Iowa II *de novo* assembly using the BWA-MEM alignment algorithm with default parameters. The resulting alignments were converted to SAM format, sorted into BAM format, and deduplicated using Picard tools. Variants for each sample were called using HaplotypeCaller and were subsequently aggregated across all samples using GenotypeGVCFs (see https://github.com/ruicatxiao/Automated_Bulk_Segregant_Analysis for detailed parameters).

We compared the genome-wide distribution of SNPs between the parental parasites and identified three regions with extremely high SNP density (>30 SNPs per kb) (Figure 1C). The regions with high SNP density include: chr 4:1,099kb – 1,106kb (7kb), chr 5: 0-12kb (12kb), and chr 6: 4kb-14kb (10kb). We excluded these regions from further bulk segregant analysis, leaving a total of 3,570 SNPs for subsequent analyses. We also identified four genomic regions with notably low SNP density (<5 SNPs per 100 kb). The regions with low SNP density include: chr 1: 390kb-700kb (310kb), chr 2: 345kb-559kb (214kb), chr 4: 199kb-530kb (331kb), and chr 7: 387kb-530kb (143kb). Similar high SNP density distribution patterns were observed in population data at the same locations^71^ (Figure 4), suggesting that these are common hypervariable regions in *C. parvum*. However, no regions with extremely low SNP density were identified in the currently published population data.

### Bulk segregant analysis

SNP loci with coverage below 30× in either of the compared pools were excluded from each bulk segregant analysis. At each variable locus, we counted reads corresponding to the genotypes of each parent and calculated allele frequencies. Iowa II allele frequencies were plotted across the genome (Figure S6), and outliers were removed using Hampel’s rule with a window size of 100 loci. Bulk segregant analysis was performed using the R package *QTLseqr*. Extreme QTLs were defined as loci with false discovery rates (FDRs, Benjamini-Hochberg adjusted *p*-values) below 0.01. Once a QTL was detected, a 95% confidence interval was estimated using Li’s method to localize causative genes^72^.

### Automated bulk segregant analysis

We also developed an Automated Bulk Segregant Analysis (ABSA) pipeline implemented in Python (v3.10.12) to streamline the workflow. This links the software needed: raw reads are quality-trimmed using Trim Galore, aligned to a reference genome with BWA, processed using SAMTools, and variants are called with GATK4 (v4.6.1.0) and filtered via vcffilter. Downstream analyses and visualization are performed in R. ABSA supports both manual installation and containerized deployment via Singularity, with a provided Singularity definition file based on Ubuntu 24.04 that bundles all required dependencies into a single, reproducible image. The code can be accessed at (https://github.com/ruicatxiao/Automated_Bulk_Segregant_Analysis).

## Supplementary Figures

**Supplementary Figure S1.**
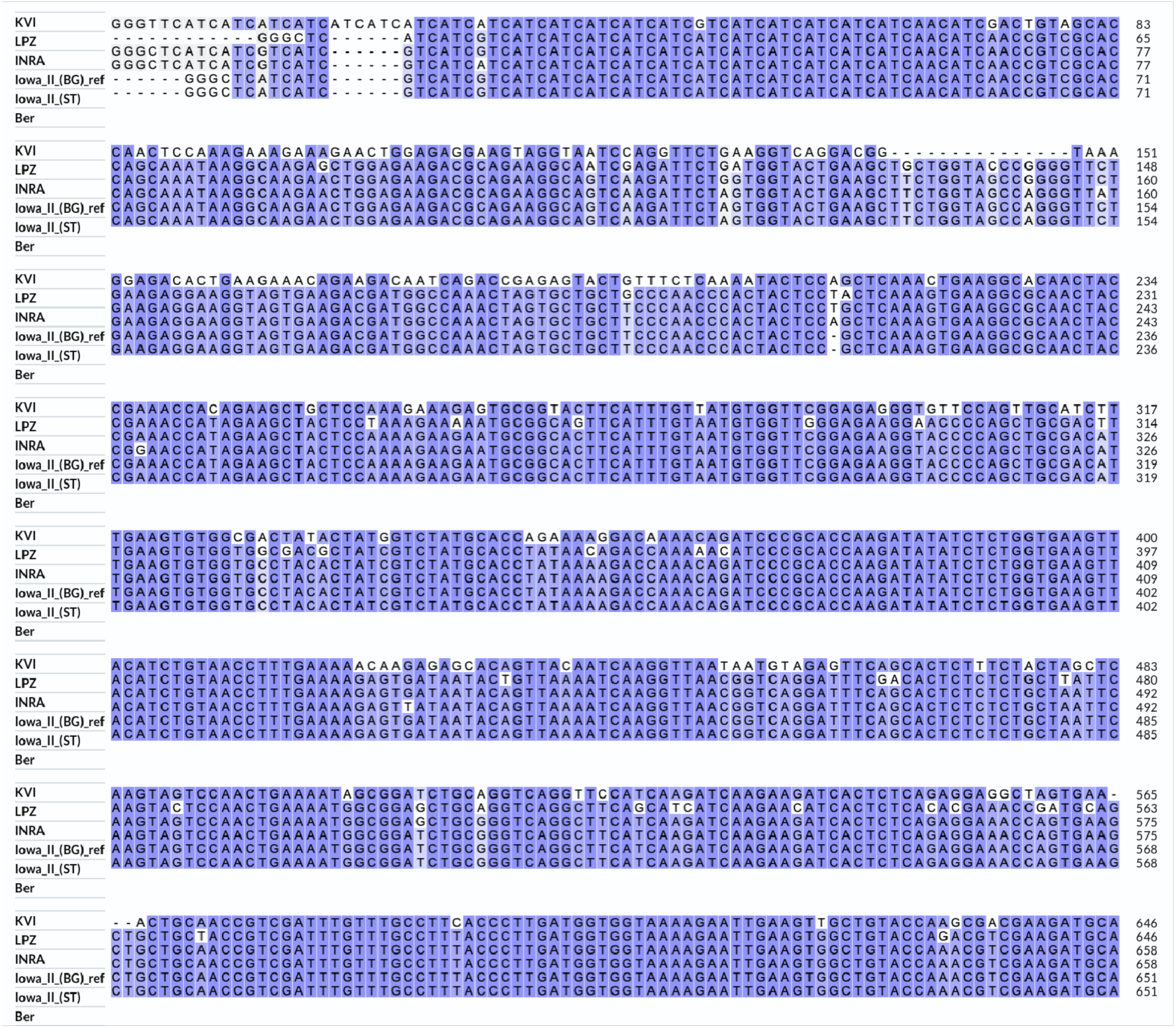
Genotyping of the strains used in this study. Multiple sequence alignment of the sequence used for GP60 genotyping of KVI, LPZ, INRA, Iowa II (ST) and Ber to the reference genome (Iowa II (BG)^25^) generated using ClustalW. Dashes represent nucleotides absent from the respective gene.

**Supplementary Figure S2.**
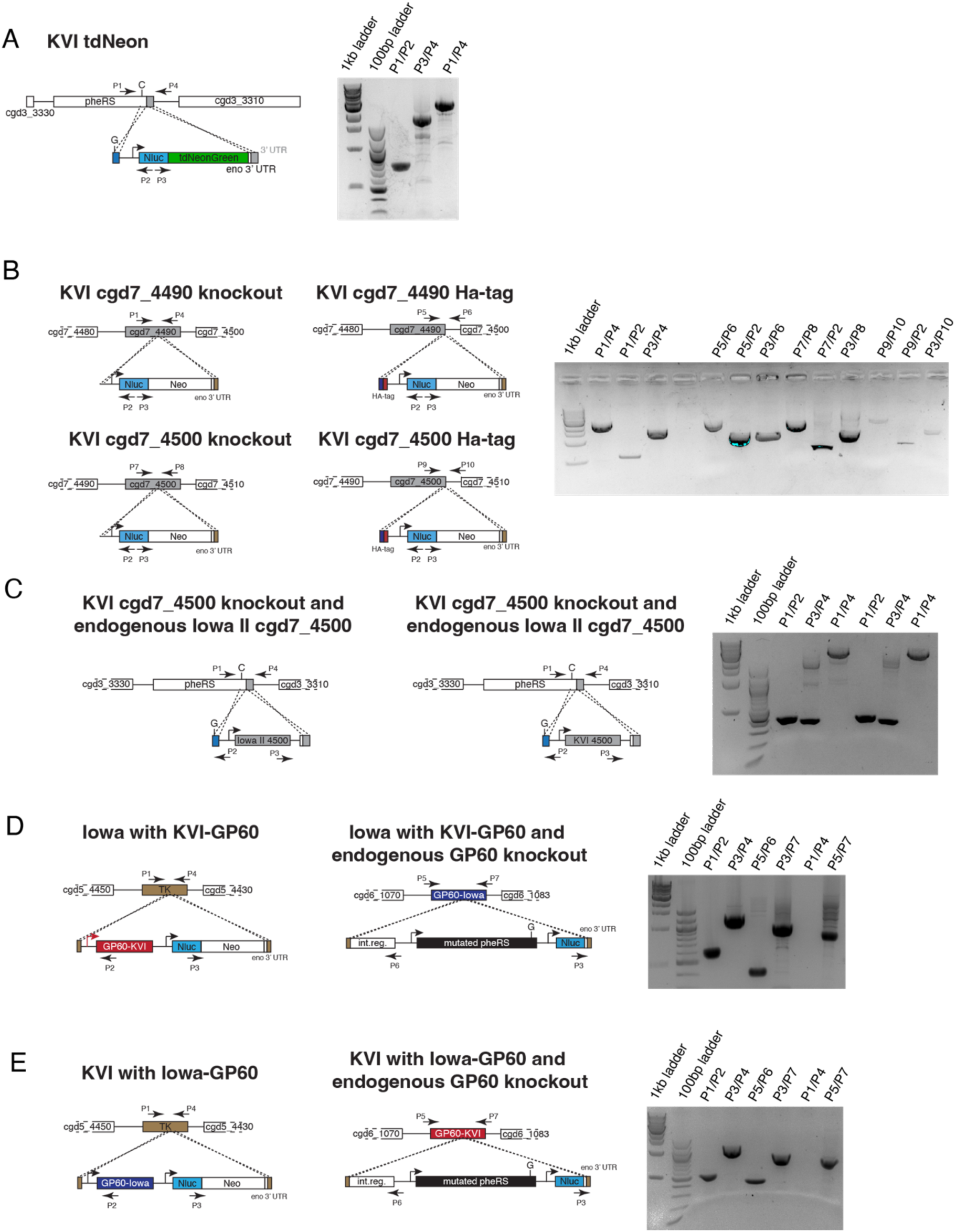
Transgenic parasite strains constructed for this study. (A) Map of the KVI tdNeon strain used in the crosses. Gel shows PCR mapping of the strain. Labelling of the gel refers to the amplicons shown in the map. (B) Knockout and in situ HA-tag mutants for the genes cgd7_4490 (ΔDG5) and cgd7_4500 (ΔDG6). Labelling of the gel refers to the amplicons shown in the maps. (C) Addback mutants (either with the cgd7_4500- Iowa II or cgd7_4500-KVI version) in cgd7_4500 knockout transgenic (ΔDG6). Labelling of the gel refers to the amplicons shown in the maps. (D) Left map shows the Iowa II strain expressing GP60-KVI in the TK locus. Right map shows the Iowa II strain with an endogenous GP60 knockout and expressing GP60-KVI in the TK locus. Labelling of the gel refers to the amplicons shown in the maps. (E) Strains generated for the allelic replacement. Left map shows the KVI strain expressing GP60-Iowa II in the TK locus. Right map shows the KVI strain with an endogenous GP60 knockout and expressing GP60-Iowa II in the TK locus. Labelling of the gel refer to the amplicons shown in the maps. A list of oligos used to genotype the strains can be found in Supplementary Table S1, under a tab sheet called ‘Genotyping’.

**Supplementary Figure S3.**
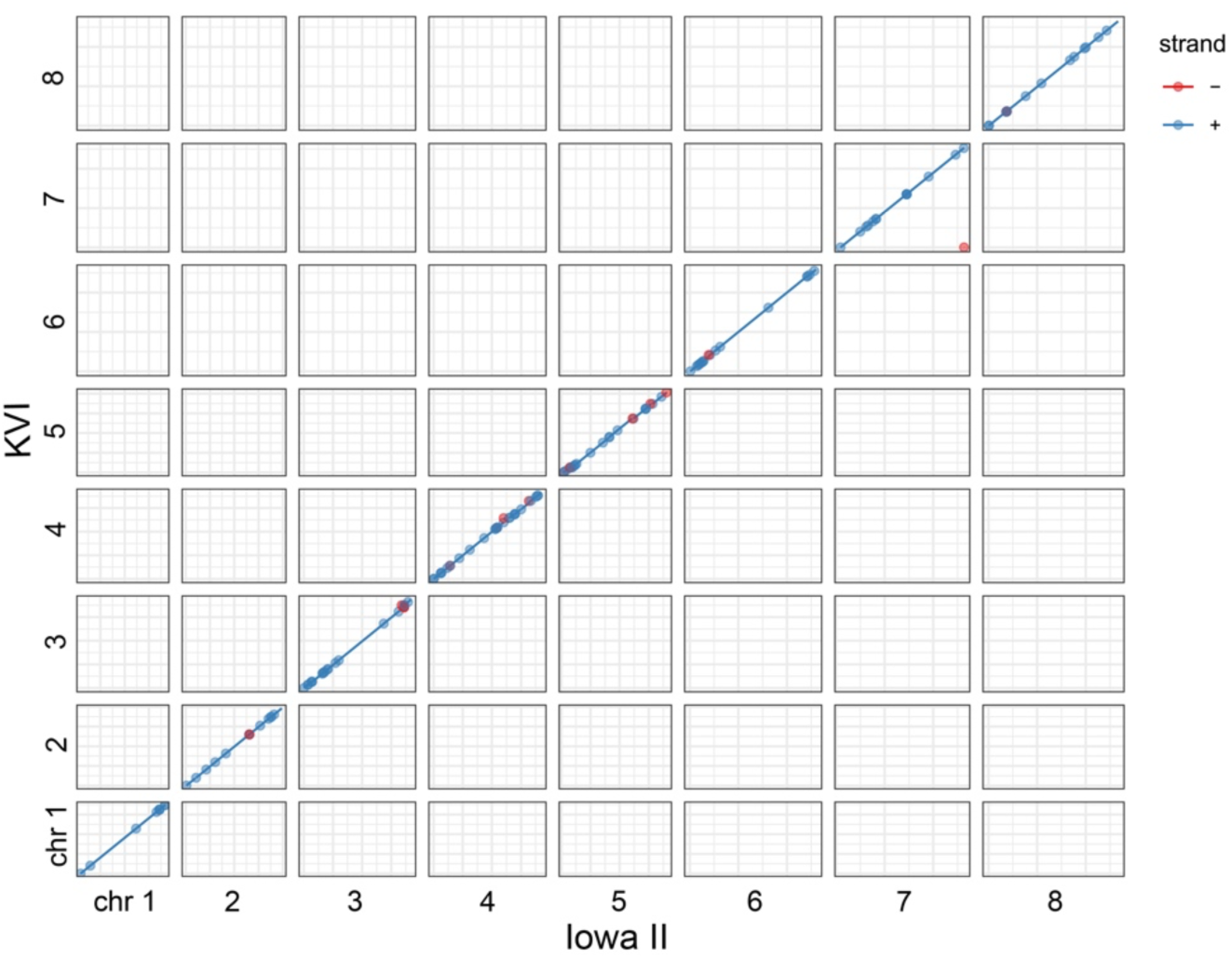
Comparison of the Iowa II and KVI genomes. The Iowa II and KVI genomes share a high degree of similarity, with only a few small inversions and one translocation throughout the genome. This translocation (∼500bp) was observed at the end of chr 7, and represents a rearrangement of the hemogen gene. The inversions ranged from 100 bp to 53 kb.

**Supplementary Figure S4.**
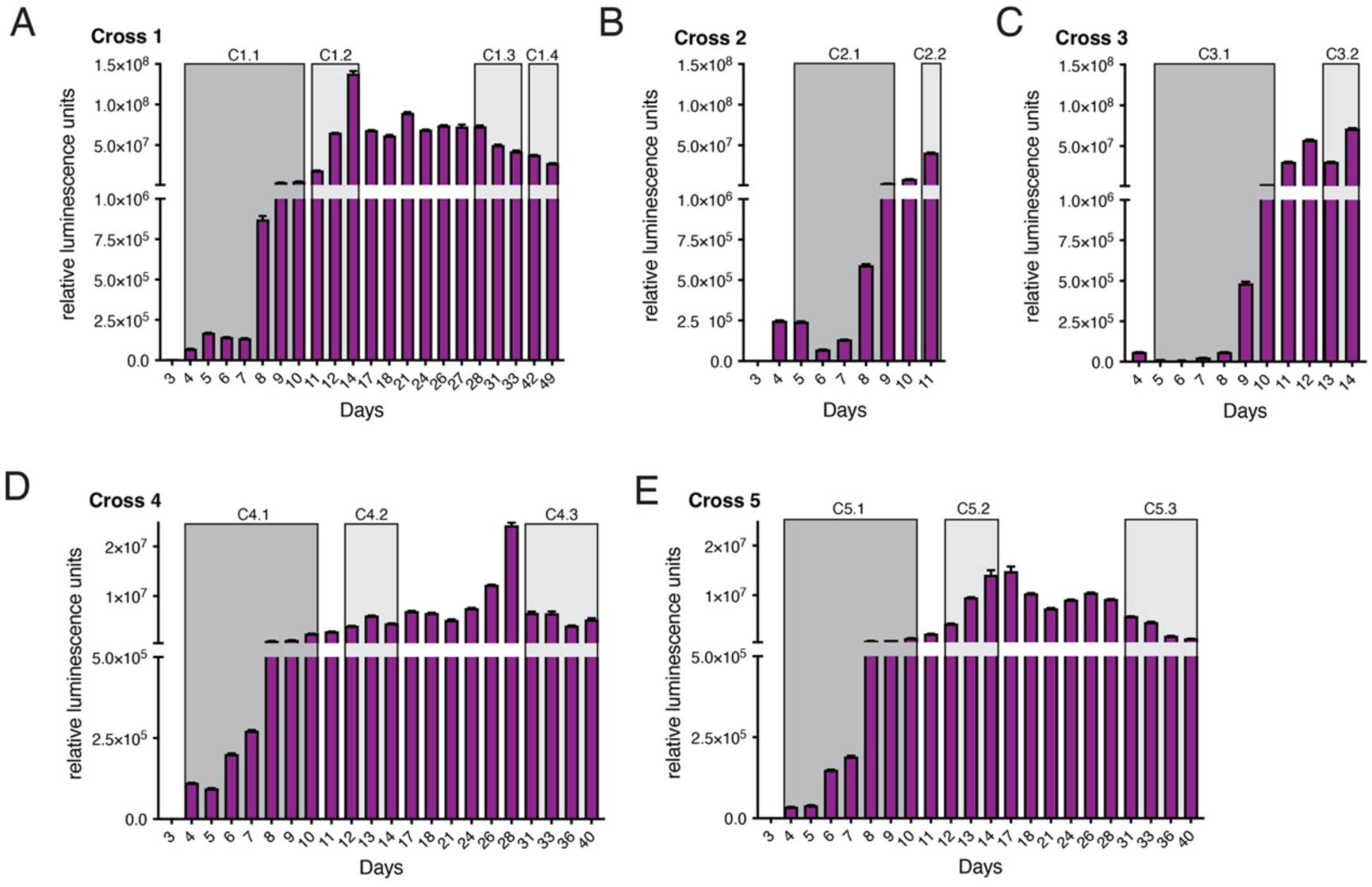
Parasite burden and sampling windows for all genetic crosses conducted in this study. Genetic crosses with (A-C) and without (D and E) dual drug treatment. Parasite burden was measured by following fecal luciferase activity. Grey boxes indicate the days for collected feces was pooled for subsequent oocyst purification and whole genome sequencing. The impact of treatment with paromomycin enhances parasite growth in mice and can lead to more severe infection. Two experiments under treatment had to be ended early, on days 11 and 14 (Figure S4B and S4C, respectively), in accordance with the endpoint defined in the animal protocol.

**Supplementary Figure S5.**
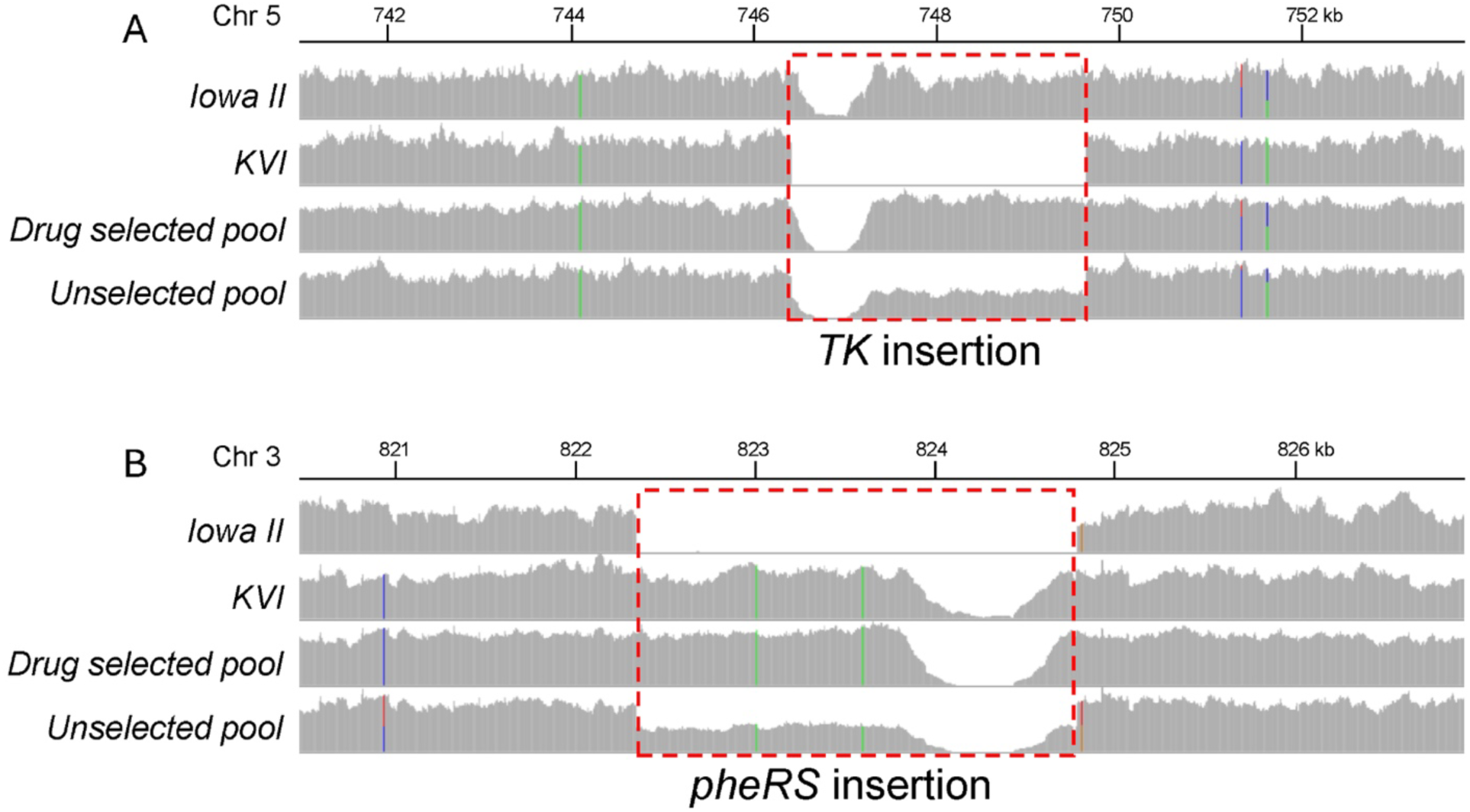
Sequence reads mapping to the resistance cassette insertion sites in parental strains and cross progeny. Read coverage plots show uniquely mapped reads for resistance cassette insertion site (A, *TK* locus; B, *pheRS* locus) and the surrounding regions. We observed complete replacement of the wildtype alleles with the resistance cassette insertions at both the *TK* and the *pheRS* locus in all drug-selected pools. The gap regions in both plots A and B corresponded to promoter sequences used in the marker construct (these sequences are naturally present in the genome, and upon transfection of the additional copy now no longer recognized as uniquely mapped and thus ignored). Note balance 50/50 representation of both alleles in the unselected pools.

**Supplementary Figure S6.**
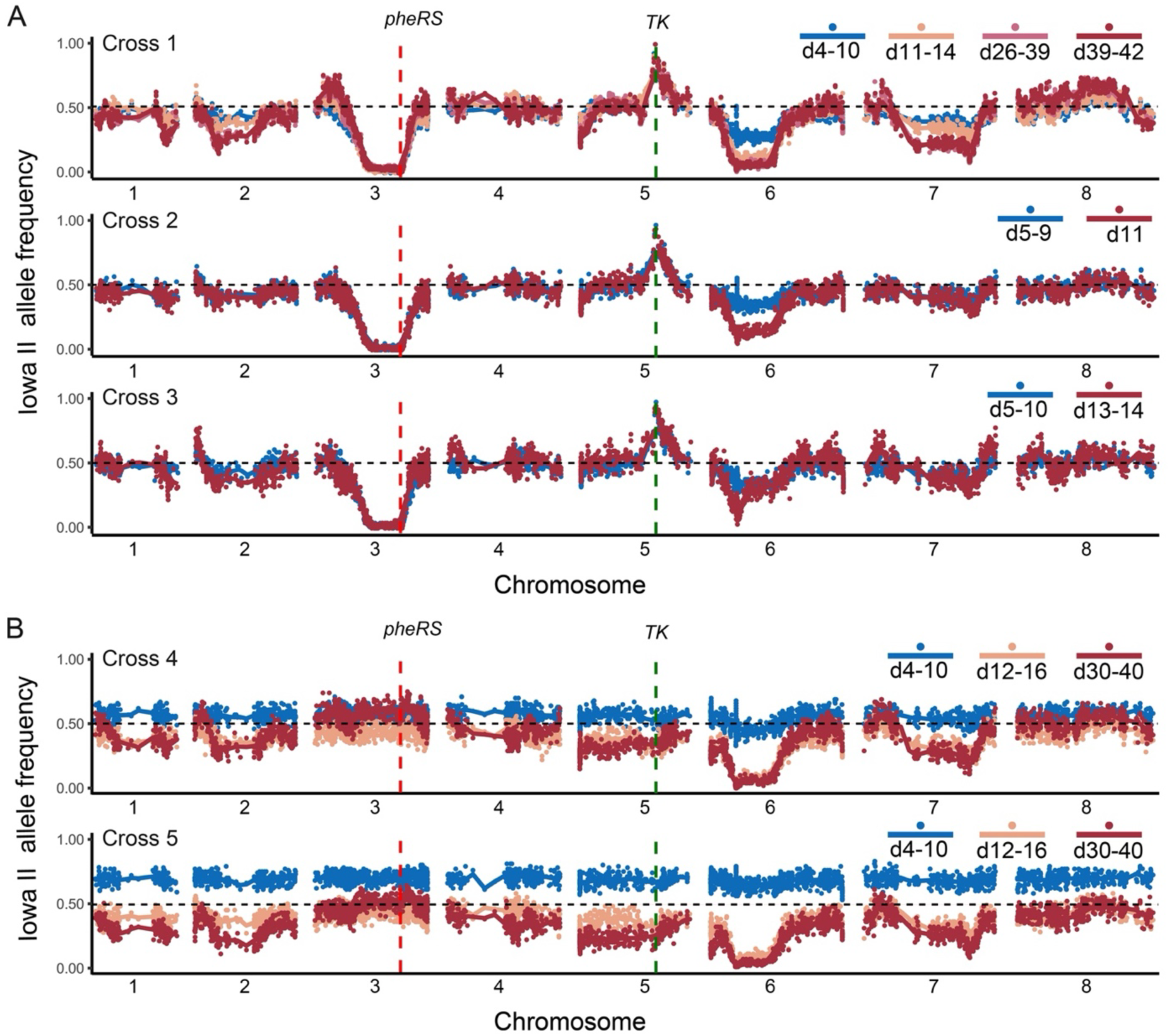
Summary of allele frequencies of all genetic crosses conducted in this study. (A) Analysis of crosses 1-3 conducted with dual BRD7929/paromomycin selection and (B) without drug selection. Allele frequencies across all chromosomes compared to the Iowa II genome (1 indicates 100% Iowa II, whereas 0 indicates 100% KVI).

**Supplementary Figure S7.**
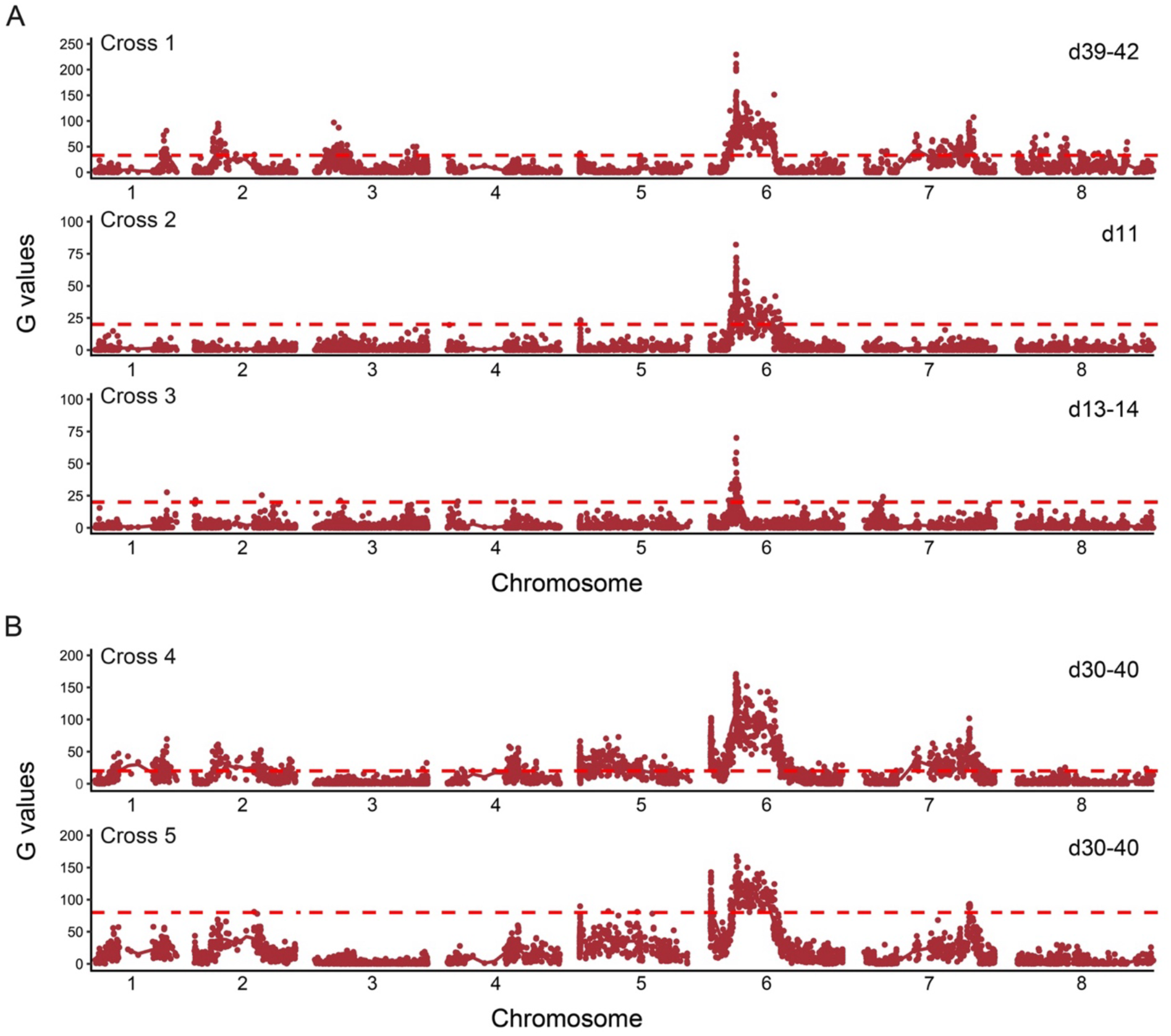
Summary of bulk segregant analysis of all genetic crosses conducted in this study. (A) Analysis of crosses 1-3 conducted with dual BRD7929/paromomycin selection and (B) without drug selection. Analysis is based on comparison of the genome sequence derived from first oocyst pool collected with the last pool (specific days varied for the last pool and the number of days are indicated for each experiment).

**Supplementary Figure S8.**
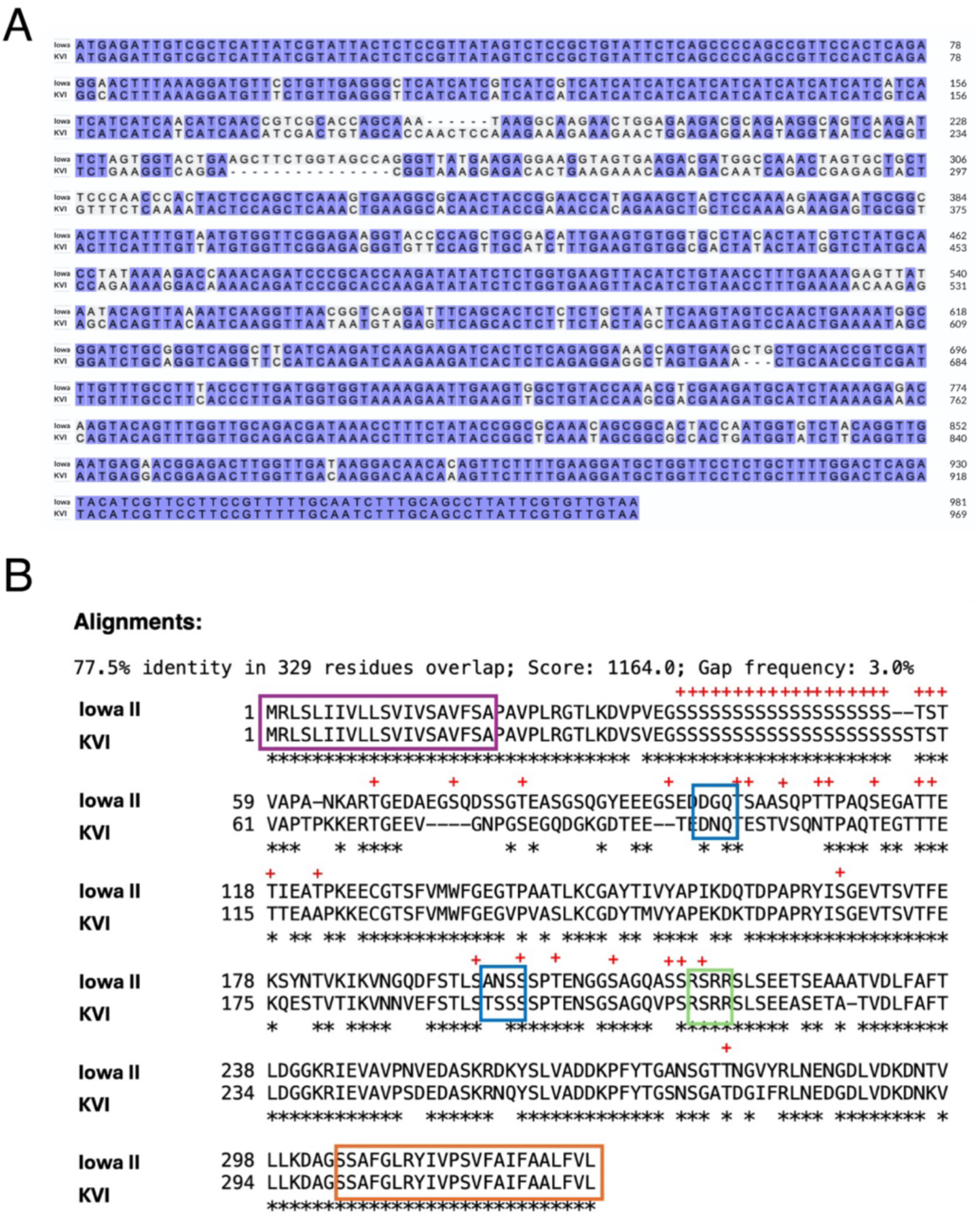
Alignment of the GP60 sequences of Iowa II and KVI. (A) A multiple sequence alignment was generated using ClustalW based on similarity. (B) The purple box represents the signal peptide, the orange box the GPI anchor, and the green box the furin cleavage site. The asterisk represents amino acids identical between the strains, while the dashes represent amino acid absent in one strain. The blue boxes show potential N-glycosylation sites in either the KVI or the Iowa II strain, and the red plus signs indicated potential O-glycosylation sites in the Iowa II strain.

## References

1. Checkley, W., White, A.C., Jr., Jaganath, D., Arrowood, M.J., Chalmers, R.M., Chen, X.M., Fayer, R., Griffiths, J.K., Guerrant, R.L., Hedstrom, L., et al. (2015). A review of the global burden, novel diagnostics, therapeutics, and vaccine targets for cryptosporidium. Lancet Infect Dis 15, 85–94. 10.1016/S1473-3099(14)70772-8.

2. Striepen, B. (2013). Parasitic infections: Time to tackle cryptosporidiosis. Nature 503, 189–191.

3. Robertson, L.J., Campbell, A.T., and Smith, H.V. (1992). Survival of Cryptosporidium parvum oocysts under various environmental pressures. Appl Environ Microbiol 58, 3494–3500. 10.1128/aem.58.11.3494-3500.1992.

4. Kotloff, K.L., Nataro, J.P., Blackwelder, W.C., Nasrin, D., Farag, T.H., Panchalingam, S., Wu, Y., Sow, S.O., Sur, D., Breiman, R.F., et al. (2013). Burden and aetiology of diarrhoeal disease in infants and young children in developing countries (the Global Enteric Multicenter Study, GEMS): a prospective, case-control study. Lancet 382, 209–222. 10.1016/S0140-6736(13)60844-2.

5. Korpe, P.S., and Petri, W.A., Jr. (2012). Environmental enteropathy: critical implications of a poorly understood condition. Trends Mol Med 18, 328–336. 10.1016/j.molmed.2012.04.007.

6. Mondal, D., Haque, R., Sack, R.B., Kirkpatrick, B.D., and Petri, W.A., Jr. (2009). Attribution of malnutrition to cause-specific diarrheal illness: evidence from a prospective study of preschool children in Mirpur, Dhaka, Bangladesh. Am J Trop Med Hyg 80, 824–826.

7. Wojcik, G.L., Korpe, P., Marie, C., Mentzer, A.J., Carstensen, T., Mychaleckyj, J., Kirkpatrick, B.D., Rich, S.S., Concannon, P., Faruque, A.S.G., et al. (2020). Genome-Wide Association Study of Cryptosporidiosis in Infants Implicates PRKCA. mBio 11. 10.1128/mBio.03343-19.

8. Kabir, M., Alam, M., Nayak, U., Arju, T., Hossain, B., Tarannum, R., Khatun, A., White, J.A., Ma, J.Z., Haque, R., et al. (2021). Nonsterile immunity to cryptosporidiosis in infants is associated with mucosal IgA against the sporozoite and protection from malnutrition. PLoS Pathog 17, e1009445. 10.1371/journal.ppat.1009445.

9. Charania, R., Wade, B.E., McNair, N.N., and Mead, J.R. (2020). Changes in the Microbiome of Cryptosporidium-Infected Mice Correlate to Differences in Susceptibility and Infection Levels. Microorganisms 8. 10.3390/microorganisms8060879.

10. Cohn, I.S., Henrickson, S.E., Striepen, B., and Hunter, C.A. (2022). Immunity to Cryptosporidium: Lessons from Acquired and Primary Immunodeficiencies. J Immunol 209, 2261–2268. 10.4049/jimmunol.2200512.

11. Pardy, R.D., Wallbank, B.A., Striepen, B., and Hunter, C.A. (2023). Immunity to Cryptosporidium: insights into principles of enteric responses to infection. Nat Rev Immunol. 10.1038/s41577-023-00932-3.

12. Sateriale, A., Slapeta, J., Baptista, R., Engiles, J.B., Gullicksrud, J.A., Herbert, G.T., Brooks, C.F., Kugler, E.M., Kissinger, J.C., Hunter, C.A., and Striepen, B. (2019). A Genetically Tractable, Natural Mouse Model of Cryptosporidiosis Offers Insights into Host Protective Immunity. Cell Host Microbe 26, 135–146 e135. 10.1016/j.chom.2019.05.006.

13. Feng, Y., Ryan, U.M., and Xiao, L. (2018). Genetic Diversity and Population Structure of Cryptosporidium. Trends Parasitol 34, 997–1011. 10.1016/j.pt.2018.07.009.

14. Jia, R., Huang, W., Huang, N., Yu, Z., Li, N., Xiao, L., Feng, Y., and Guo, Y. (2022). High infectivity and unique genomic sequence characteristics of Cryptosporidium parvum in China. PLoS neglected tropical diseases 16, e0010714. 10.1371/journal.pntd.0010714.

15. Huang, W., Guo, Y., Lysen, C., Wang, Y., Tang, K., Seabolt, M.H., Yang, F., Cebelinski, E., Gonzalez-Moreno, O., Hou, T., et al. (2023). Multiple introductions and recombination events underlie the emergence of a hyper-transmissible Cryptosporidium hominis subtype in the USA. Cell Host Microbe 31, 112–123 e114. 10.1016/j.chom.2022.11.013.

16. Slapeta, J. (2013). Cryptosporidiosis and Cryptosporidium species in animals and humans: a thirty colour rainbow? Int J Parasitol 43, 957–970. 10.1016/j.ijpara.2013.07.005.

17. English, E.D., Guerin, A., Tandel, J., and Striepen, B. (2022). Live imaging of the Cryptosporidium parvum life cycle reveals direct development of male and female gametes from type I meronts. PLoS Biol 20, e3001604. 10.1371/journal.pbio.3001604.

18. Tandel, J., English, E.D., Sateriale, A., Gullicksrud, J.A., Beiting, D.P., Sullivan, M.C., Pinkston, B., and Striepen, B. (2019). Life cycle progression and sexual development of the apicomplexan parasite Cryptosporidium parvum. Nat Microbiol 4, 2226–2236. 10.1038/s41564-019-0539-x.

19. Walzer, K.A., Tandel, J., Byerly, J.H., Daniels, A.M., Gullicksrud, J.A., Whelan, E.C., Carro, S.D., Krespan, E., Beiting, D.P., and Striepen, B. (2024). Transcriptional control of the Cryptosporidium life cycle. Nature. 10.1038/s41586-024-07466-1.

20. Kimball, A., Funkhouser-Jones, L., Huang, W., Xu, R., Witola, W.H., and Sibley, L.D. (2024). Mendelian segregation and high recombination rates facilitate genetic analyses in Cryptosporidium parvum. PLoS genetics 20, e1011162. 10.1371/journal.pgen.1011162.

21. Behnke, M.S., Dubey, J.P., and Sibley, L.D. (2016). Genetic Mapping of Pathogenesis Determinants in Toxoplasma gondii. Annu Rev Microbiol 70, 63–81. 10.1146/annurev-micro-091014-104353.

22. Su, X., Hayton, K., and Wellems, T.E. (2007). Genetic linkage and association analyses for trait mapping in Plasmodium falciparum. Nat Rev Genet 8, 497–506. 10.1038/nrg2126.

23. Shaw, S., Cohn, I.S., Baptista, R.P., Xia, G., Melillo, B., Agyabeng-Dadzie, F., Kissinger, J.C., and Striepen, B. (2024). Genetic crosses within and between species of Cryptosporidium. Proc Natl Acad Sci U S A 121, e2313210120. 10.1073/pnas.2313210120.

24. Tako, S., Fleiderovitz, L., Markovich, M.P., Mazuz, M.L., Behar, A., and Yasur-Landau, D. (2023). Cryptosporidium parvum gp60 subtypes in diarrheic lambs and goat kids from Israel. Parasitol Res 122, 2237–2241. 10.1007/s00436-023-07925-0.

25. de Paula Baptista, R., Xiao, R., Li, Y., Glenn, T.C., and Kissinger, J.C. (2023). New T2T assembly of Cryptosporidium parvum IOWA annotated with reference genome gene identifiers. bioRxiv. 10.1101/2023.06.13.544219.

26. Robinson, G., Chalmers, R.M., Elwin, K., Guy, R.A., Bessonov, K., Troell, K., and Xiao, L. (2025). Deciphering a cryptic minefield: A guide to Cryptosporidium gp60 subtyping. Curr Res Parasitol Vector Borne Dis 7, 100257. 10.1016/j.crpvbd.2025.100257.

27. Vinayak, S., Pawlowic, M.C., Sateriale, A., Brooks, C.F., Studstill, C.J., Bar-Peled, Y., Cipriano, M.J., and Striepen, B. (2015). Genetic modification of the diarrhoeal pathogen Cryptosporidium parvum. Nature 523, 477–480. 10.1038/nature14651.

28. Li, X., Kumar, S., Brenneman, K.V., and Anderson, T.J.C. (2022). Bulk segregant linkage mapping for rodent and human malaria parasites. Parasitology international 91, 102653. 10.1016/j.parint.2022.102653.

29. Guerin, A., Strelau, K.M., Barylyuk, K., Wallbank, B.A., Berry, L., Crook, O.M., Lilley, K.S., Waller, R.F., and Striepen, B. (2023). Cryptosporidium uses multiple distinct secretory organelles to interact with and modify its host cell. Cell Host Microbe 31, 650–664 e656. 10.1016/j.chom.2023.03.001.

30. Strong, W.B., Gut, J., and Nelson, R.G. (2000). Cloning and sequence analysis of a highly polymorphic Cryptosporidium parvum gene encoding a 60-kilodalton glycoprotein and characterization of its 15- and 45-kilodalton zoite surface antigen products. Infect Immun 68, 4117–4134.

31. Bouzid, M., Hunter, P.R., Chalmers, R.M., and Tyler, K.M. (2013). Cryptosporidium pathogenicity and virulence. Clin Microbiol Rev 26, 115–134. 10.1128/CMR.00076-12.

32. Li, M., Yang, F., Hou, T., Gong, X., Li, N., Sibley, L.D., Feng, Y., Xiao, L., and Guo, Y. (2024). Variant surface protein GP60 contributes to host infectivity of Cryptosporidium parvum. Commun Biol 7, 1175. 10.1038/s42003-024-06885-0.

33. Taylor, S., Barragan, A., Su, C., Fux, B., Fentress, S.J., Tang, K., Beatty, W.L., Hajj, H.E., Jerome, M., Behnke, M.S., et al. (2006). A secreted serine-threonine kinase determines virulence in the eukaryotic pathogen Toxoplasma gondii. Science 314, 1776–1780. 10.1126/science.1133643.

34. Saeij, J.P., Boyle, J.P., Coller, S., Taylor, S., Sibley, L.D., Brooke-Powell, E.T., Ajioka, J.W., and Boothroyd, J.C. (2006). Polymorphic secreted kinases are key virulence factors in toxoplasmosis. Science 314, 1780–1783. 10.1126/science.1133690.

35. Jex, A.R., and Tyler, K.M. (2023). American variants of Cryptosporidium hominis: Over-sexed and over here? Cell Host Microbe 31, 5–7. 10.1016/j.chom.2022.12.012.

36. Tichkule, S., Jex, A.R., van Oosterhout, C., Sannella, A.R., Krumkamp, R., Aldrich, C., Maiga-Ascofare, O., Dekker, D., Lamshoft, M., Mbwana, J., et al. (2021). Comparative genomics revealed adaptive admixture in Cryptosporidium hominis in Africa. Microb Genom 7. 10.1099/mgen.0.000493.

37. Nader, J.L., Mathers, T.C., Ward, B.J., Pachebat, J.A., Swain, M.T., Robinson, G., Chalmers, R.M., Hunter, P.R., van Oosterhout, C., and Tyler, K.M. (2019). Evolutionary genomics of anthroponosis in Cryptosporidium. Nat Microbiol 4, 826–836. 10.1038/s41564-019-0377-x.

38. Corsi, G.I., Tichkule, S., Sannella, A.R., Vatta, P., Asnicar, F., Segata, N., Jex, A.R., van Oosterhout, C., and Caccio, S.M. (2023). Recent genetic exchanges and admixture shape the genome and population structure of the zoonotic pathogen Cryptosporidium parvum. Mol Ecol 32, 2633–2645. 10.1111/mec.16556.

39. Talenti, A., Powell, J., Hemmink, J.D., Cook, E.A.J., Wragg, D., Jayaraman, S., Paxton, E., Ezeasor, C., Obishakin, E.T., Agusi, E.R., et al. (2022). A cattle graph genome incorporating global breed diversity. Nature communications 13, 910. 10.1038/s41467-022-28605-0.

40. Feng, Y., Li, N., Roellig, D.M., Kelley, A., Liu, G., Amer, S., Tang, K., Zhang, L., and Xiao, L. (2017). Comparative genomic analysis of the IId subtype family of Cryptosporidium parvum. Int J Parasitol 47, 281–290. 10.1016/j.ijpara.2016.12.002.

41. Chantepie, S., and Chevin, L.M. (2020). How does the strength of selection influence genetic correlations? Evol Lett 4, 468–478. 10.1002/evl3.201.

42. Vinayak, S., Jumani, R.S., Miller, P., Hasan, M.M., McLeod, B.I., Tandel, J., Stebbins, E.E., Teixeira, J.E., Borrel, J., Gonse, A., et al. (2020). Bicyclic azetidines kill the diarrheal pathogen Cryptosporidium in mice by inhibiting parasite phenylalanyl-tRNA synthetase. Sci Transl Med 12. 10.1126/scitranslmed.aba8412.

43. Braun, L., Brenier-Pinchart, M.P., Hammoudi, P.M., Cannella, D., Kieffer-Jaquinod, S., Vollaire, J., Josserand, V., Touquet, B., Coute, Y., Tardieux, I., et al. (2019). The Toxoplasma effector TEEGR promotes parasite persistence by modulating NF-kappaB signalling via EZH2. Nat Microbiol 4, 1208–1220. 10.1038/s41564-019-0431-8.

44. Hakimi, M.A., Olias, P., and Sibley, L.D. (2017). Toxoplasma Effectors Targeting Host Signaling and Transcription. Clinical Microbiology Reviews 30, 615–645. 10.1128/Cmr.00005-17.

45. Olias, P., Etheridge, R.D., Zhang, Y., Holtzman, M.J., and Sibley, L.D. (2016). Toxoplasma Effector Recruits the Mi-2/NuRD Complex to Repress STAT1 Transcription and Block IFN-gamma-Dependent Gene Expression. Cell Host Microbe 20, 72–82. 10.1016/j.chom.2016.06.006.

46. Laurent, F., and Lacroix-Lamande, S. (2017). Innate immune responses play a key role in controlling infection of the intestinal epithelium by Cryptosporidium. International Journal for Parasitology 47, 711–721. 10.1016/j.ijpara.2017.08.001.

47. Brockhurst, M.A., Chapman, T., King, K.C., Mank, J.E., Paterson, S., and Hurst, G.D. (2014). Running with the Red Queen: the role of biotic conflicts in evolution. Proc Biol Sci 281. 10.1098/rspb.2014.1382.

48. O’Connor, R.M., Wanyiri, J.W., Cevallos, A.M., Priest, J.W., and Ward, H.D. (2007). Cryptosporidium parvum glycoprotein gp40 localizes to the sporozoite surface by association with gp15. Mol Biochem Parasit 156, 80–83. 10.1016/j.molbiopara.2007.07.010.

49. Cevallos, A.M., Zhang, X., Waldor, M.K., Jaison, S., Zhou, X., Tzipori, S., Neutra, M.R., and Ward, H.D. (2000). Molecular cloning and expression of a gene encoding Cryptosporidium parvum glycoproteins gp40 and gp15. Infect Immun 68, 4108–4116.

50. Cavailles, P., Flori, P., Papapietro, O., Bisanz, C., Lagrange, D., Pilloux, L., Massera, C., Cristinelli, S., Jublot, D., Bastien, O., et al. (2014). A highly conserved Toxo1 haplotype directs resistance to toxoplasmosis and its associated caspase-1 dependent killing of parasite and host macrophage. PLoS Pathog 10, e1004005. 10.1371/journal.ppat.1004005.

51. McNair, N.N., Bedi, C., Shayakhmetov, D.M., Arrowood, M.J., and Mead, J.R. (2018). Inflammasome components caspase-1 and adaptor protein apoptosis-associated speck-like proteins are important in resistance to Cryptosporidium parvum. Microbes Infect 20, 369–375. 10.1016/j.micinf.2018.04.006.

52. Sateriale, A., Gullicksrud, J.A., Engiles, J.B., McLeod, B.I., Kugler, E.M., Henao-Mejia, J., Zhou, T., Ring, A.M., Brodsky, I.E., Hunter, C.A., and Striepen, B. (2021). The intestinal parasite Cryptosporidium is controlled by an enterocyte intrinsic inflammasome that depends on NLRP6. Proc Natl Acad Sci U S A 118. 10.1073/pnas.2007807118.

53. Gibson, A.R., Sateriale, A., Dumaine, J.E., Engiles, J.B., Pardy, R.D., Gullicksrud, J.A., O’Dea, K.M., Doench, J.G., Beiting, D.P., Hunter, C.A., and Striepen, B. (2022). A genetic screen identifies a protective type III interferon response to Cryptosporidium that requires TLR3 dependent recognition. PLoS Pathog 18, e1010003. 10.1371/journal.ppat.1010003.

54. O’Hara, S.P., Bogert, P.S., Trussoni, C.E., Chen, X., and LaRusso, N.F. (2011). TLR4 promotes Cryptosporidium parvum clearance in a mouse model of biliary cryptosporidiosis. J Parasitol 97, 813–821. 10.1645/GE-2703.1.

55. Mead, J.R. (2023). Early immune and host cell responses to Cryptosporidium infection. Front Parasitol 2. 10.3389/fpara.2023.1113950.

56. Wang, Y., Gao, M., Li, X., Zhu, W., Zhao, M., Li, J., Liu, X., Cao, L., Li, S., Zhang, S., et al. (2023). Structural Analyses of a Dominant Cryptosporidium parvum Epitope Presented by H-2K(b) Offer New Options To Combat Cryptosporidiosis. mBio 14, e0266622. 10.1128/mbio.02666-22.

57. Bhalchandra, S., Gevers, K., Heimburg-Molinaro, J., van Roosmalen, M., Coppens, I., Cummings, R.D., and Ward, H.D. (2023). Identification of the glycopeptide epitope recognized by a protective Cryptosporidium monoclonal antibody. Infect Immun 91, e0027523. 10.1128/iai.00275-23.

58. Martinelli, A., Cheesman, S., Hunt, P., Culleton, R., Raza, A., Mackinnon, M., and Carter, R. (2005). A genetic approach to the de novo identification of targets of strain-specific immunity in malaria parasites. Proc Natl Acad Sci U S A 102, 814–819. 10.1073/pnas.0405097102.

59. Lacroix-Lamande, S., Mancassola, R., Naciri, M., and Laurent, F. (2002). Role of gamma interferon in chemokine expression in the ileum of mice and in a murine intestinal epithelial cell line after Cryptosporidium parvum infection. Infect Immun 70, 2090–2099. 10.1128/IAI.70.4.2090-2099.2002.

60. Xiao, L.H., Morgan, U.M., Limor, J., Escalante, A., Arrowood, M., Shulaw, W., Thompson, R.C.A., Fayer, R., and Lal, A.A. (1999). Genetic diversity within Cryptosporidium parvum and related Cryptosporidium species. Appl Environ Microb 65, 3386–3391.

61. Xiao, L., Alderisio, K., Limor, J., Royer, M., and Lal, A.A. (2000). Identification of species and sources of Cryptosporidium oocysts in storm waters with a small-subunit rRNA-based diagnostic and genotyping tool. Appl Environ Microbiol 66, 5492–5498. 10.1128/AEM.66.12.5492-5498.2000.

62. Feng, Y., Li, N., Duan, L., and Xiao, L. (2009). Cryptosporidium genotype and subtype distribution in raw wastewater in Shanghai, China: evidence for possible unique Cryptosporidium hominis transmission. J Clin Microbiol 47, 153–157. 10.1128/JCM.01777-08.

63. Altschul, S.F., Madden, T.L., Schaffer, A.A., Zhang, J., Zhang, Z., Miller, W., and Lipman, D.J. (1997). Gapped BLAST and PSI-BLAST: a new generation of protein database search programs. Nucleic Acids Res 25, 3389–3402.

64. Alves, L.M., Heisler, E.G., Kissinger, J.C., Patterson, J.M., and Kalan, E.B. (1979). Effects of Controlled Atmospheres on Production of Sesquiterpenoid Stress Metabolites by White Potato Tuber: Possible Involvement of Cyanide-resistant Respiration. Plant Physiol 63, 359–362. 10.1104/pp.63.2.359.

65. 65. Cryptosporidiosis. (2022). In WOAH Terrestrial Manual.

66. Kato, S., Jenkins, M.B., Ghiorse, W.C., and Bowman, D.D. (2001). Chemical and physical factors affecting the excystation of Cryptosporidium parvum oocysts. J Parasitol 87, 575–581. 10.1645/0022-3395(2001)087[0575:CAPFAT]2.0.CO;2.

67. Pawlowic, M.C., Vinayak, S., Sateriale, A., Brooks, C.F., and Striepen, B. (2017). Generating and Maintaining Transgenic Cryptosporidium parvum Parasites. Curr Protoc Microbiol 46, 20B 22 21-20B 22 32. 10.1002/cpmc.33.

68. Dos Santos Pacheco, N., and Soldati-Favre, D. (2021). Coupling Auxin-Inducible Degron System with Ultrastructure Expansion Microscopy to Accelerate the Discovery of Gene Function in Toxoplasma gondii. Methods Mol Biol 2369, 121–137. 10.1007/978-1-0716-1681-9_8.

69. Liffner, B., Cepeda Diaz, A.K., Blauwkamp, J., Anaguano, D., Frolich, S., Muralidharan, V., Wilson, D.W., Dvorin, J.D., and Absalon, S. (2023). Atlas of Plasmodium falciparum intraerythrocytic development using expansion microscopy. Elife 12. 10.7554/eLife.88088.

70. Mansfield, B.E., Oltean, H.N., Oliver, B.G., Hoot, S.J., Leyde, S.E., Hedstrom, L., and White, T.C. (2010). Azole Drugs Are Imported By Facilitated Diffusion in Candida albicans and Other Pathogenic Fungi. Plos Pathogens 6. ARTN e1001126 10.1371/journal.ppat.1001126.

